# Slowly-conducting pyramidal tract neurons in macaque and rat

**DOI:** 10.1101/768770

**Authors:** A Kraskov, D Soteropoulos, I Glover, RN Lemon, SN Baker

## Abstract

Anatomical studies report a large proportion of fine myelinated fibres in the primate pyramidal tract (PT), while very few pyramidal tract neurons (PTNs) with slow conduction velocities (CV) (< ∼10 m/s) are reported electrophysiologically. This discrepancy might reflect recording bias towards fast PTNs or prevention of antidromic invasion by recurrent inhibition of slow PTNs from faster axons. We investigated these factors in recordings made with a polyprobe (32 closely-spaced contacts) from motor cortex of anaesthetised rats (n=2) and macaques (n=3), concentrating our search on PTNs with long antidromic latencies. We identified 21 rat PTNs with antidromic latencies > 2.6 ms and estimated CV 3-8 m/s, and 67 macaque PTNs (> 3.9ms, CV 6-12 m/s). Spikes of most slow PTNs were small and present on only some recording contacts, while spikes from simultaneously recorded fast-conducting PTNs were large and appeared on all contacts. Antidromic thresholds were similar for fast and slow PTNS, while spike duration was considerably longer in slow PTNs. Most slow PTNs showed no signs of failure to respond antidromically. A number of tests, including intracortical microinjection of bicuculline (GABA_A_ antagonist), failed to provide any evidence that recurrent inhibition prevented antidromic invasion of slow PTNs. Our results suggest that recording bias is the main reason why previous studies were dominated by fast PTNs.

## Introduction

The corticospinal tract is known to subserve a number of different functions (Kuypers 1981; Lemon 2008) and this is probably reflected in the wide range of fibre diameters present in the tract, particularly in primates (Häggqvist 1937; Lassek 1941; Firmin et al. 2014). Pyramidal tract neurons (PTNs) can be antidromically activated from the PT, and studying the activity of these neurons during a wide range of motor and other tasks has been a particularly fruitful approach to understanding corticospinal function (Evarts 1965; Turner and Delong, 2000; Kraskov et al. 2009; Vigneswaran et al. 2011; Quallo et al. 2012). However, anatomical and electrophysiological approaches to the primate corticospinal tract reveal very contrasting accounts. While anatomical studies have emphasized the huge preponderance of fine, myelinated axons within the tract, with axon diameters of 0.5-3 µm (Häggqvist 1937; Russell and Demyer 1961; Innocenti et al. 2019), recordings of antidromic responses evoked in PTNs by stimulation of the PT are dominated by responses of neurons with relatively short antidromic latencies (ADLs), indicating large, fast conducting axons (e.g. (Humphrey and Corrie 1978; Firmin et al. 2014). Recordings from primary motor cortex (M1), premotor and supplementary motor cortex (Evarts 1965; Humphrey and Corrie 1978; Macpherson et al. 1982; Firmin et al. 2014) all reveal a strong bias toward fast-conducting PTNs.

Responses from ‘slow PTNs’, with conduction velocities < 10 m/s, which, on anatomical grounds, might be expected to make up the bulk of the responses, are largely missing from published studies. As a result, our understanding of how the different components of the corticospinal system function is far from complete. A first step in understanding the role of the slow-conducting neurons would be to discover why slow PTNs are so under-represented in electrophysiological studies. Once methods for identifying these neurons have been developed, it should be possible to study the functional contribution of these slow PTNs in awake animals, using the same approaches that have been so successful in the study of the fast-conducting PTNs.

There are a number of possible explanations for the ‘corticospinal discrepancy’ between anatomical and electrophysiological studies (Kraskov et al. 2019). Firstly, it is possible that conventional stimulation parameters applied to the PT fail to excite the finer axons within the tract. Secondly, slow PTNs might be expected to arise from neurons with small cell bodies and these might be under-represented in extracellular recording studies, because of the well-known bias in such recording towards large neurons (Towe and Harding 1970; Humphrey and Corrie 1978). A final possibility is inhibition or collision resulting from recurrent synaptic actions on slow PTNs caused by stimulation of fast corticospinal axons. These recurrent effects would act upon slow PTNs well before antidromic impulses in their own axons arrive at the cell body. Recurrent inhibition could hyperpolarise slow PTNs and prevent antidromic invasion, as suggested by Innocenti et al. (2019).

In this study we tested a silicon probe with multiple (32) recording sites spaced at high density on a single shaft. We used these probes in a series of experiments in anaesthetised rats and macaque monkeys. The first experiments, in the rat, were a good test of the electrode being capable of recording from slow PTNs, because the population of PTNs in the rat is relatively homogenous, with small pyramidal neurons having axons of up to 2.5 µm in diameter; rodents lack the large, fast PTNs seen in primates and other species (Mediratta and Nicoll 1983; Leenen et al. 1985). These experiments established that we could antidromically identify PTNs in rat sensorimotor cortex with latencies likely to reflect conduction velocities <10 m/s, and that PT stimulation intensities used were in a range similar to that used for antidromic activation of fast PTNs in the macaque. Subsequently, we used these same approaches in the macaques to search in primary motor cortex for slow PTNs with long ADLs, again corresponding to conduction velocities <10 m/s. We purposely focused on PTNs with longer ADLs. We found no evidence that recurrent inhibition from fast PTNs prevented antidromic invasion of slow PTNs.

## Materials and Methods

All experiments were approved by the Animal Welfare and Ethical Review Board of Newcastle University, where they were carried out, and performed under appropriate personal and project licences issued by the UK Home Office.

### Preparatory surgery

#### Macaque monkeys

Three adult female Rhesus macaques (L, N and O, body weights: 5.9, 6.9 and 8.8 kg, respectively), which were all purpose bred for research, were used. Macaques were socially-housed in groups. Anaesthesia was induced with ketamine (10 mg/kg i.m.), supplemented by medetomidine (3 μg/kg i.m.) and midazolam (0.3 mg/kg i.m.) (monkey O) or propofol (1.3-4.4 mg/kg, monkeys L and N) General anaesthesia during surgery consisted of sevoflurane (2-3% in 100% oxygen) and alfentanil (0.4-0.57 μg/kg/min i.v.). Meloxicam (0.3 mg/kg i.m., all animals) and paracetamol (25 mg/kg, monkey O only) were given to augment analgesia. Antibiotic was given to reduce the risk of infection (cefotaxime 20 mg/kg i.v. or co-amoxiclav, 12.5-20 mg/kg i.v.). Methylprednisolone (5.4 mg/kg/hr i.v.) was given to reduce cerebral oedema. Hartmann’s solution was given to prevent dehydration (approximately 5 ml/kg/hr; rates adjusted to provide, with drug infusions, a total rate of around 10 ml/kg/hr). A tracheotomy was made, allowing positive pressure artificial ventilation. A central arterial line inserted via the carotid artery provided continual arterial blood pressure monitoring. The bladder was catheterised. The animal’s temperature was maintained by a warm air blanket and thermostatic heat pad.

After placing in a stereotaxic head holder, a craniotomy was made over the right motor cortex and the dura removed. A stimulating electrode (tungsten insulated with parylene, Microprobes Inc, part number LF501G) with a tip impedance of ∼ 10 kΩ was advanced into the medullary PT on the right side, via a craniotomy extending rostrally from the foramen magnum, at an angle of 30° and with coordinates 2.0 mm caudal to obex and 1.0 mm from the midline. The antidromic volley from the surface of the ipsilateral (right) motor cortex was recorded, and the electrode fixed at the point of lowest threshold (8-9 mm below obex, volley threshold <25 µA). A laminectomy was made to expose the cervical spinal cord and stimulating electrodes, identical to those in the PT, were then implanted into the lateral funiculus on the left side at C3 and at C6 around 1.5-2.0 mm below the surface of the cord.

After all surgery was completed, anaesthesia was switched to continuous i.v. infusion of alfentanil (0.4-1.2 μg/kg/hr), ketamine (6-10 mg/kg/hr) and midazolam (0.3 mg/kg/hr), and sevoflurane was reduced (to 0% in monkeys L, N; to 0.5% in monkey O), as we have found that this regime provides good central nervous system activity whilst maintaining a stable plane of anaesthesia. Anaesthetic monitoring throughout consisted of pulse oximetry, heart rate, arterial blood pressure, core and peripheral temperature, end-tidal CO_2_. Rapid increases of heart rate or blood pressure in response to noxious stimuli, or slowly increasing trends in either measure, were taken as indicative of waning levels of anaesthesia and supplementary doses of the injectable agents were given.

#### Rats

Two adult male Sprague-Dawley rats (body weights: 345g and 360g) were anaesthetised with urethane (1.3 g/kg i.p.) and buprenorphine (10 µg/kg i.p.), supplemented with isoflurane (1-2% in 100% oxygen). All surgery was carried out under additional isoflurane. The head was mounted in a stereotaxic head holder. A craniotomy was made over the left sensorimotor cortex and the dura removed. As in the monkeys, the brainstem was exposed by a craniotomy rostral to the foramen magnum, and a single stimulating electrode advanced into the left pyramidal tract (the electrode was positioned 1 mm rostral and 0.5 mm lateral to obex), while monitoring the antidromic volley from the ipsilateral motor cortex. The electrode was fixed around 4.5 mm below obex with a volley threshold of ∼25 µA.

### Recording and stimulation

A 32-contact silicon recording probe (poly3, NeuroNexus) was mounted on a piezoelectric micromanipulator (Newport part number PZA12), and angled to penetrate the primary motor cortex normal to the cortical surface. The probe (shaft width 114 μm, 15 μm thick) had 32 contacts (diameter 15 μm; typical impedance at 1 kHz 750 kΩ) arranged in an array which spanned a region 290 μm high by 51 μm wide. The probe was connected to a digital headstage (Intan technologies, RHD2132; bandpass 1 Hz-10 kHz, gain 200) and digital data were sampled to computer disc at 25 kSamples/s (Intan Technologies Recording Controller). The probe was slowly advanced into the cortex (2 µm step every 1.5 s) using a modified version of a program previously developed to locate intracellular recordings with micropipette electrodes automatically (Collins and Baker 2014). Critically, this program synchronised search stimuli applied to the PT with the recording electrode movement cycle, so that sweeps were not corrupted by movement artefact.

The PT search stimuli consisted of two shocks delivered 10 ms apart (intensity ≥750 µA; charge-balanced biphasic stimuli, 100 µs per phase). This helped to identify putative antidromic responses, since these almost always followed both stimuli, whereas synaptic responses were usually far less consistent to the second shock. In addition, while the response to the first stimulus could show some jitter in latency, probably related to previous spontaneous activity in the neuron, responses to the second stimulus showed less jitter (Swadlow et al. 1978).

Another advantage of this double shock was to test for effects of recurrent inhibition blocking antidromic invasion. Since early work had suggested that recurrent inhibition from a PT shock might take several milliseconds to build up (Stefanis and Jasper 1964; Takahashi et al. 1967) the second shock, at 10 ms, would have arrived when inhibition should be well-developed. Therefore, we predicted that if inhibition blocked antidromic invasion of a PTN, this should result in a lack of responses in the PTN to the second shock.

The experimenter stopped the automated electrode advance as soon as a putative antidromic response was identified, making subsequent small adjustments via manual commands to the piezoelectric manipulator as required to optimise the recording.

### Identification of antidromic responses in slow PTNs

In each track we searched for neurons showing responses at a relatively fixed latency after the PT stimulus. There was only a small amount of jitter in the ADL, presumably reflecting periods of super- and subnormal activity in the axon after passage of a spontaneous spike (Swadlow et al. 1978). Stimulation rate was 0.5 Hz. We deliberately ignored short-latency responses from fast PTNs and instead searched for PTNs with latencies of >2.5 ms (rat) or >3.8 ms (macaque). After determining the threshold (T) for each cell, all further tests, including latency measurements, were carried out at 1.2xT. A number of criteria were used to confirm responses as antidromic (see Results). We checked that responses had low jitter in latency, a sharp and stable threshold, and followed a train of high frequency stimuli (three shocks, inter-stimulus interval (ISI) 3 ms). ‘Perfect’ following (a response to every stimulus in the train on every sweep (up to 300 sweeps were checked) was found in the majority of slow PTNs; however, a minority showed failures on some sweeps. Some of the tested neurons were spontaneously active and this allowed us to confirm collision of the antidromic responses (Lemon 1984).

### Further tests to characterise slow PTNs

In both rats and macaques we looked at two additional properties of slow PTNs:

#### Responses to strong stimulation of the PT

we reasoned that if slow PTNs receive recurrent synaptic effects via collaterals of faster PT axons, these effects would be greatest when we tested maximal PT shocks (up to 2 mA). We therefore compared antidromic invasion of slow PTNs at 1.2T and then at the higher intensity.

#### Volatile anaesthesia

we reasoned that deepening anaesthesia would suppress activity in cortical neurons, blocking recurrent synaptic effects whilst leaving antidromic conduction in corticospinal axons intact. We therefore initially searched for PTNs while injectable anaesthesia was supplemented with volatile anaesthetic (isoflurane in rats, servoflurane in macaques, both at 1%). As volatile agents are known to hyperpolarise cortical cells, this should have reduced recurrent inhibition by making the interneurons less excitable. Once a slow PTN had been isolated, we turned off the gas anaesthesia for several minutes until end-tidal concentration of the volatile agent had decreased to <0.5%, before re-testing the responses to PT stimulation to determine if there was any evidence of antidromic failure. We then restored gas anaesthesia, before searching for the next PTN.

In macaques we looked at two further properties of slow PTNS:

#### Spinal termination of slow PTNs

in the macaque most PTNs have axons which continue into the spinal cord (Humphrey and Corrie 1978). In two macaques (L,N), we tested each PTN for antidromic responses from stimulating electrodes in the rostral (C3 level) and caudal (C6) dorsolateral funiculus (DLF) of the cervical cord. Search stimuli up to 2 mA were used.

#### Intracortical injection of bicuculline

in the final part of two experiments (macaques N and O), the GABA_A_ antagonist, bicuculline, was microinjected close to the recording probe. Bicuculline should block recurrent inhibition generated by PT stimulation, relieving any blockade of antidromic invasion and thereby unmasking antidromic responses in slow PTNs. After the recording probe had been positioned close to slow PTNs in layer V, a fine needle (30 gauge) connected to a Hamilton syringe and automated pump (UMP3, World Precision Instruments), was lowered to a depth of 2.0-2.5 mm below the surface and a small volume (0.5-5 µL) of bicuculline methiodide (Sigma Aldrich catalogue number 14343 made up in sterile saline), was injected over a 2-5 minute period. Recording was continued for around 20-40 minutes post-injection, before further slow PTNs were sought by moving the recording probe or making new penetrations.

## Results

We recorded from 21 slow PTNs in two rats and 67 slow PTNs in three macaque monkeys.

### Recordings in Rats

The rat recordings allowed us establish that our stimulation and recording methods were capable of identifying slow-conducting PTNs. In these recordings, we were able to identify 21 slow PTNs in 7 penetrations into the forelimb area of M1. PTNs were recorded when the probe tip was 1.1 to 1.7 mm (average 1.4 mm) below the cortical surface.

An example is shown in Fig. 1A-C. This PTN had an antidromic latency of 14.6 ms, the longest we recorded in this study. Nevertheless it had a low antidromic threshold of only 75 µA. Figure 1A shows the response to a 90 µA (1.2T) shock; the antidromic response (marked •) showed very little latency jitter. Figure 1B shows the response to two PT shocks with a 10 ms ISI, which was our standard search stimulus. The PTN followed both shocks (•), with slightly less jitter in the response to the second shock. Further confirmation of the antidromic nature of the response was obtained by showing that it followed a high frequency train of 3 shocks at ISI 3 ms (Fig. 1C, •).

**Figure 1.**
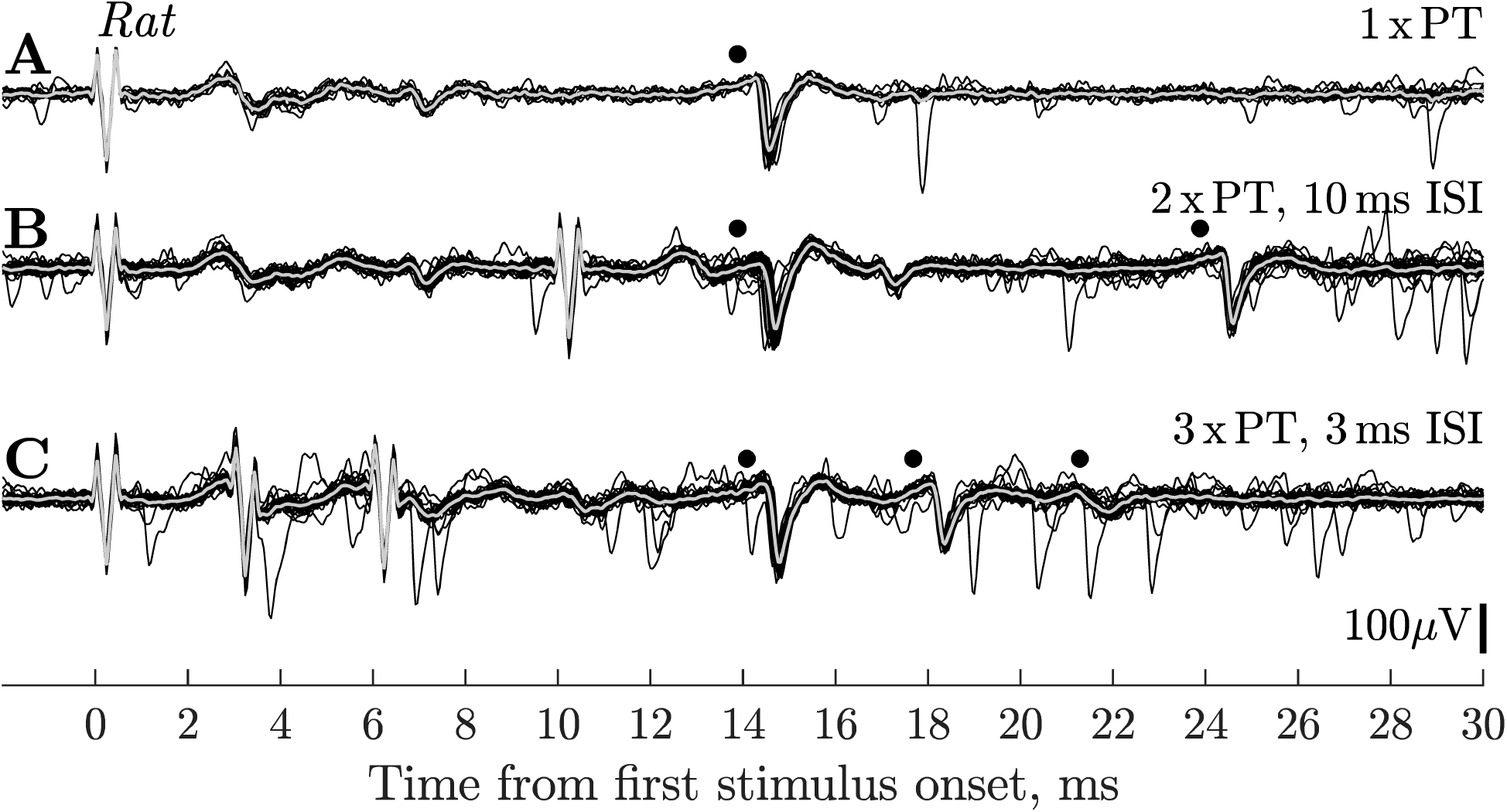
Rat. Antidromic responses of slow PTN in motor cortex. **A.** Single PT shock (90 µA) evoked a response from a slow PTN (•) at 14.6 ms. **B.** Responses to 2 x PT (90 µA) shocks with an ISI of 10ms; the PTN showed antidromic responses to both shocks (•), the amplitude of the second spike was reduced. **C.** shows responses to 3 x PT (90 uA) shocks with an ISI of 3 ms; responses (•) were present to all 3 shocks, but with a marked decrease in spike amplitude. Isoflurane anaesthesia was on for A-C. The reduction in amplitude with successive shocks was not seen in all rat PTNs, and it is unlikely that it was related to any failure in antidromic invasion, but rather was caused by changes in the spike generation mechanism when the PTNs were activated at high frequency. It is noticeable that the attenuation was less for the ISI 10 ms stimuli (B) than for ISI 3 ms (C).

We also tested whether the antidromic responses of slow PTNs with relatively low thresholds (< 120 μA) persisted when much stronger PT stimuli (up to 2mA) were applied. In all 4 PTNs tested (including the example PTN in Fig. 1), antidromic responses to both the first and second shocks were unaffected by these strong stimuli. Since such stimuli should have recruited a large amount of recurrent inhibition (see Methods), this result suggests that this inhibition was not strong enough to block antidromic invasion.

### Anaesthesia

We tried to unmask any recurrent inhibition by removing supplementary isoflurane anaesthesia, but again failed to see any differences in the yield of antidromic responses with or without isoflurane. Removal of isoflurane anaesthesia increased the level of spontaneous activity but failed to reveal any new slow PTNs in the recordings. This was tested for 14 slow PTNs.

### Latency and threshold of antidromic responses

Figure 2A shows the ADLs of 17 slow rat PTNs which could be measured accurately without contamination from other units or field potentials. ADLs were measured from the first clear inflection in the antidromic spike in averages of responses to a single PT shock. Most of these PTNs had ADLs between 4 and 8 ms. The corresponding thresholds are shown in Fig. 2B; although a few PTNs had very high thresholds, most were below 300 µA. There was no significant correlation between ADL and threshold for this population (N=21, r=0.06, p>0.5).

**Figure 2.**
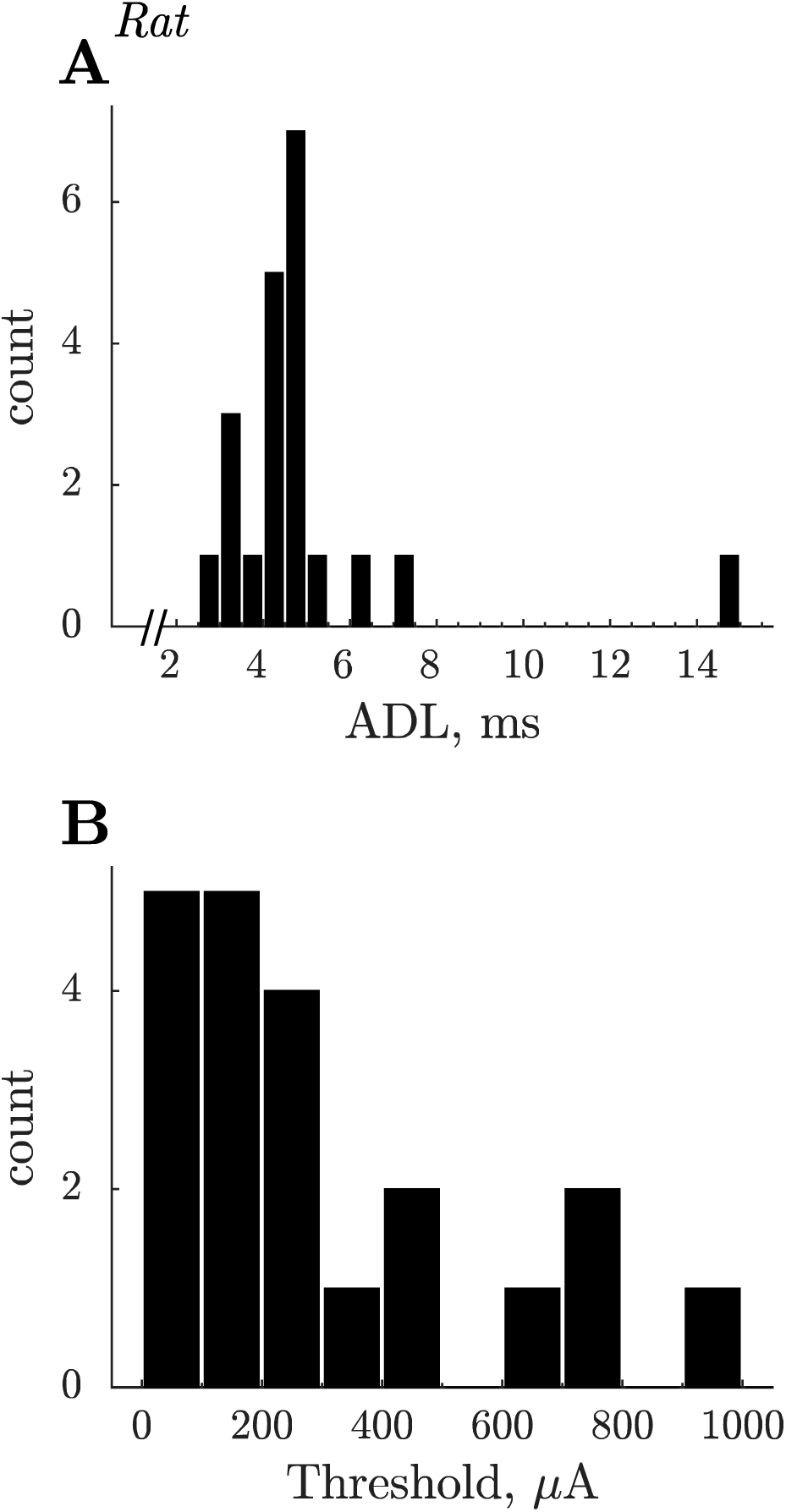
Rat. Summary of slow PTN properties. **A.** Antidromic latencies of 21 slow PTNs. Latencies were measured to the onset of the antidromic spike evoked by PT shocks with intensity of 1.2xT. The latency range of these PTNs would correspond to conduction velocities of 8 m/s down to 1.4 m/s, assuming a PT-M1 conduction distance of 20 mm and a utilisation time of 0.1 ms. **B.** Distribution of current thresholds for antidromic responses.

### Recordings in Macaques

The rat experiments demonstrated the capacity of the multiple contact silicone probe to record from PTNs with slowly-conducting axons. We therefore proceeded to use the same methodology in 3 macaque experiments. The yield of slow PTNs was 67 PTNs in 25 successful penetrations (13 in 8 penetrations in monkey L; 15 in 8 in monkey N; 39 in 9 in monkey O). The PTNs were recorded from the arm/hand area of M1 in penetrations made into the convexity of the precentral gyrus with the probe tip 0.9 to 1.8 mm (average 1.3 mm) below the cortical surface. The yield of PTNs varied substantially from one penetration to the next. The best recordings in terms of signal-to-noise were generally obtained on recording sites close to the probe’s tip. Figure 3 shows that using the silicon probe it was possible to record from clusters of PTNs at a single site (black dots), including one PTN with a long ADL (3.9 ms).

**Figure 3.**
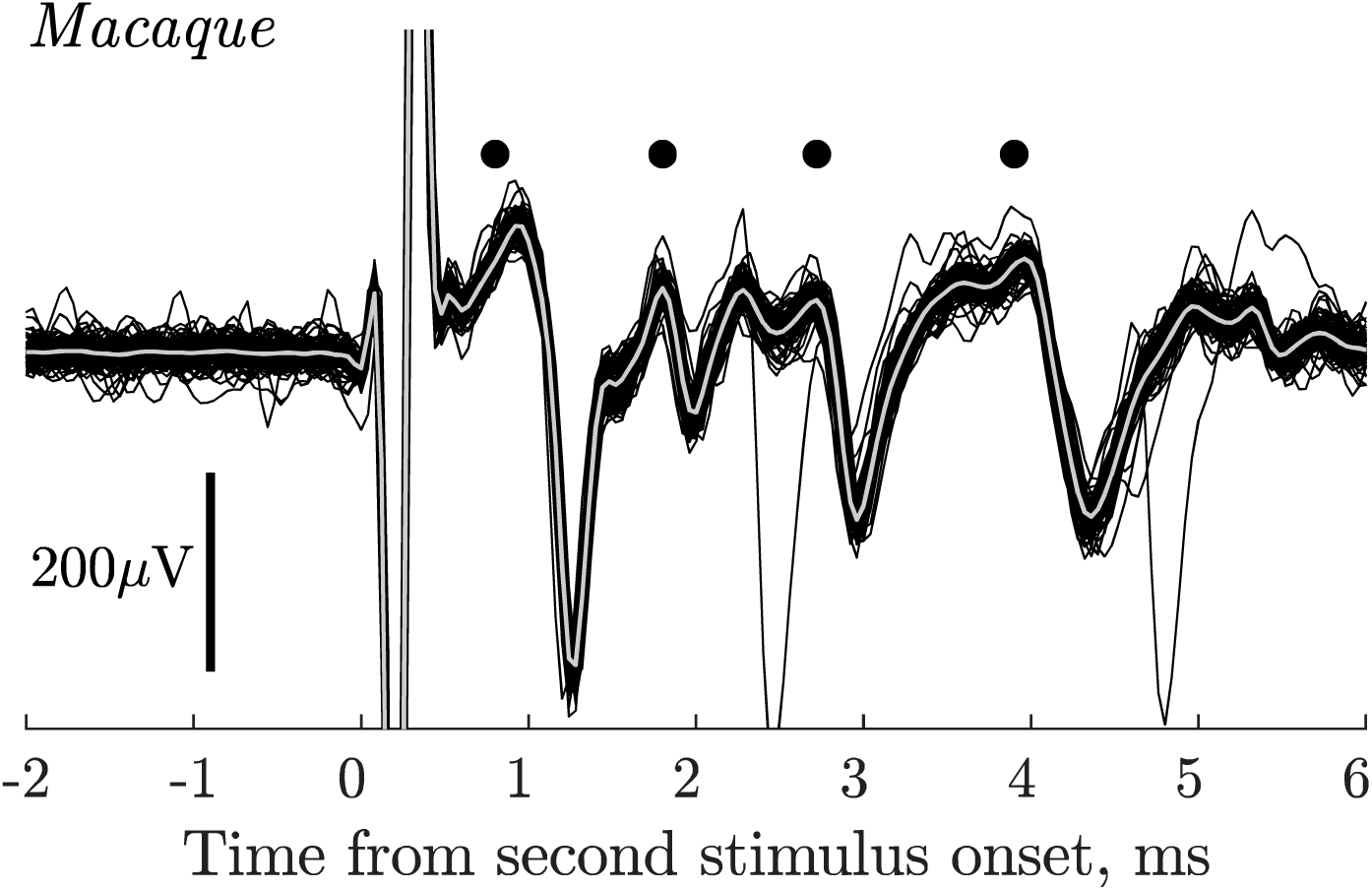
Macaque. Group of PTNs in motor cortex. Superimposed responses (100 sweeps) recorded from the polyprobe at a depth of 1.3 mm in response to PT stimulation at 100 µA intensity. In this case the standard double shock, with an ISI of 10 ms, was given (see Methods) and the responses shown are to the second shock. Antidromic responses from four PTNs are present (black dots) including one with an antidromic latency of 3.9 ms.

Figure 4A shows responses from a pair of slow PTNs with ADLs of 3.9 ms (small, earlier spike, open circle) and 5.0 ms (larger, later spike, black circle) to a single PT shock. Note that both PTNs exhibited very little jitter in antidromic latency. These PTNs were both antidromically excited from the spinal cord via an electrode in the DLF at the C3 level (Fig. 5B). The greater separation of the two spikes when activated from the cord compared with from the PT suggests variation in conduction velocity along the course of the axon.

**Figure 4.**
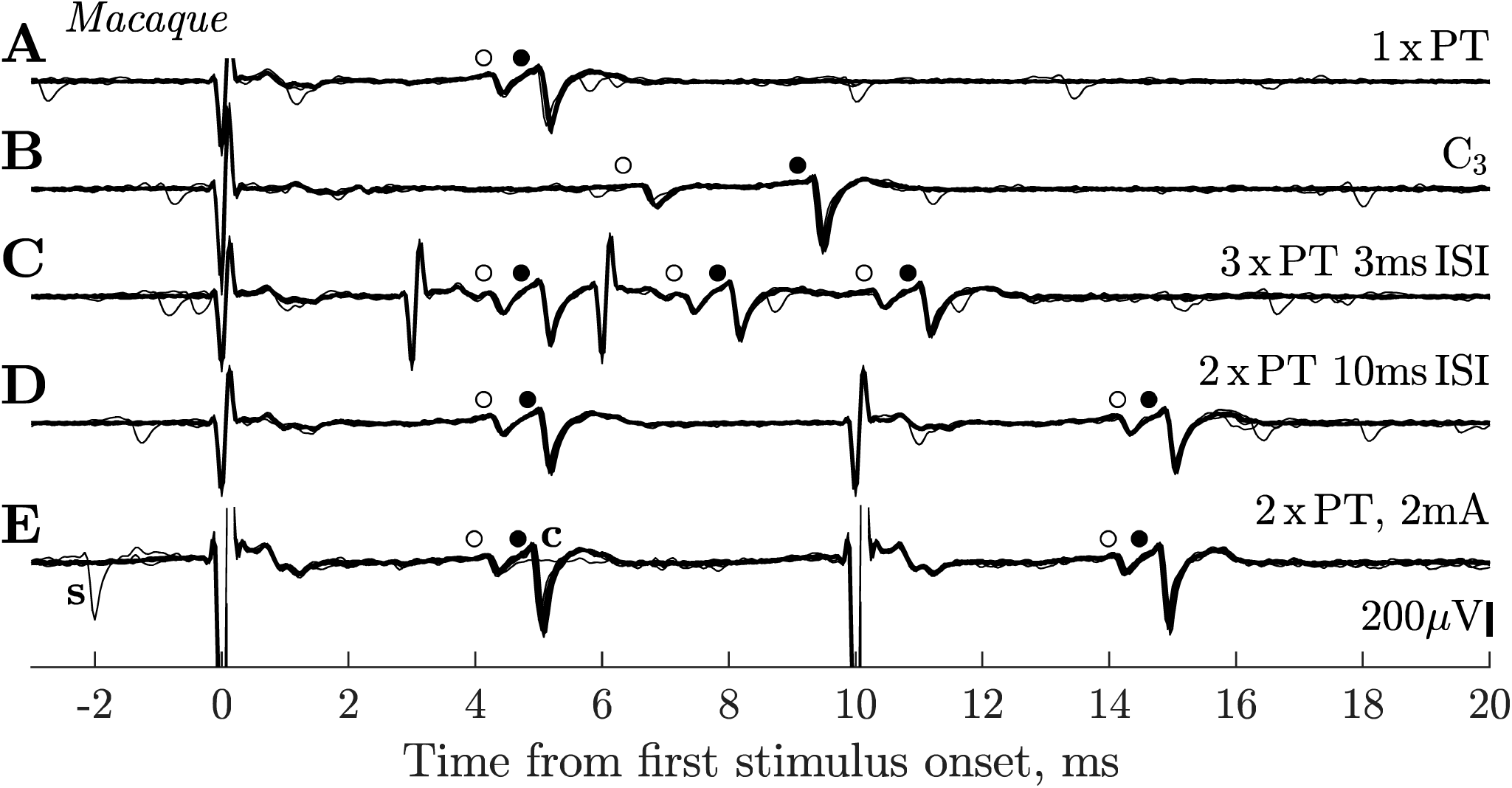
Macaque. Responses of slow PTNs to antidromic stimulation. **A**. Response of two slow PTNs to a single PT shock at 420 µA. The earlier (smaller) PTN had an ADL of 3.9 ms (o) and a threshold of 80 µA, while the later (larger) one had an ADL of 5.0 ms (•) and threshold 350 µA. **B**. These two PTNs also responded to stimulation of the spinal cord at C3 with a shock of 850 µA. Note the greater temporal separation of the two PTNs in response to the spinal vs PT stimulation. **C**. Both PTNs followed 3 x PT with an ISI of 3 ms (intensity 420 µA). All three responses had a latency and amplitude identical to that seen with a single shock. **D.** Both PTNs followed 2 x PT with an ISI of 10 ms (intensity 420 µA). **E.** Response of the same pair of PTNs to 2 x PT at high intensity (2mA). Both PTNs followed the double shock. In one sweep, a spontaneous discharge (‘s’) of the larger PTN occurred just before the first shock which resulted in collision of the larger PTN to the first shock (flat line in recording labelled ‘c’). Each row is a superimposition of 30 sweeps. Sevoflurane anaesthesia was on for A-D but off for E.

**Figure 5.**
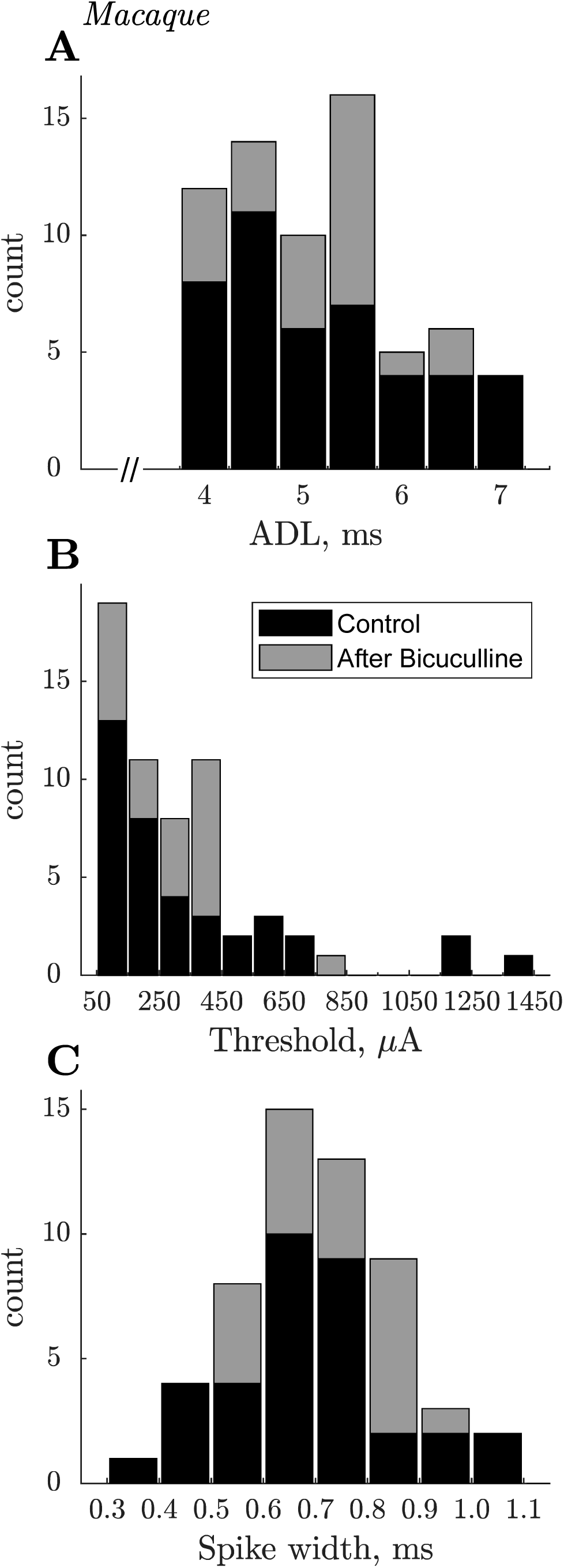
Macaque. Summary of slow PTN properties. **A.** Stacked histograms of antidromic latencies of 67 slow PTNs recorded in 3 macaques. Latencies were measured to the onset of the antidromic spike evoked by PT shocks with intensity of 1.2xT. Black and grey bars indicate PTNs recorded, respectively, before and after intracortical biccuculine. Each bin is 0.5 ms wide and represents ADLs 0.25 before, and 0.25 after, the central tick-mark (i.e. for 4 ms it was ADLs falling between 3.75 ms and 4.25 ms). The latency range of these PTNs would correspond to conduction velocities of ∼12 to ∼6 m/s, assuming a PT-M1 conduction distance of 47 mm and a utilisation time of 0.1 ms. **B.** Distribution of current thresholds for antidromic responses. Thresholds were determined in 61/67 cases. C. Distribution of spike duration in 55 slow PTNs. Spike duration was measured from the negative trough to the positive peak of each average antidromic spike. Note that nearly all slow PTNs had broad spikes with durations > 500 µS.

### High-frequency following of antidromic responses

To confirm the antidromic nature of the measured responses, we checked that each PTN could follow a high-frequency train of 3 x PT shocks with an ISI of 3 ms. Both of the PTNs in Fig. 4 showed consistent, stable responses to all three shocks (Fig. 4C). Of the 67 PTNs recorded, we were able to test 57 with high frequency 3 x PT shocks. All of them showed responses that followed all three shocks, although a minority of cases failed to exhibit ‘perfect’ following i.e. a response to all three shocks on every sweep (see below).

Figure 4D shows that both PTNs still responded to the second shock in a pair separated by an interval of 10 ms, when recurrent inhibition should be well developed. Of the 67 PTNs, we were able to test the PT double shock in 57 cases. All 57 PTNs responded to both shocks, although again a few showed less than perfect following, with the PTN occasionally failing to respond to the second shock (see below).

Figure 4E demonstrates two further points. First, in spontaneously active PTNs we were able to confirm the antidromic nature of PTN responses by demonstrating collision: a spontaneous discharge of the larger PTN, that occurred just before PT stimulation (labelled ‘s’ in Fig. 4E), collided the antidromic response to the first shock (labelled ‘c’). Second, we showed that PTNs with low thresholds (such as the smaller PTN in Fig. 4, threshold 80 µA) continued to respond even after much stronger stimuli (2 mA in Fig. 4E) were applied to the PT. Such high intensity shocks would be expected to excite a large number of fast PT fibres and generate a strong recurrent inhibitory effect. The maintained response at high intensities (which we confirmed in all 41 PTNs in which this test was carried out) suggests that recurrent inhibition exerted little effect on antidromic responses. High intensity stimulation did not shorten the ADL.

### Anaesthesia

In two monkeys (L and N), for each of 19 PTNs recorded, we tried to unmask any recurrent inhibition by temporarily lightening the anaesthesia (removing the supplementary sevoflurane), but we did not observe any differences in the yield of antidromic responses. We saw no differences in antidromic effects recorded with (e.g. Fig. 4D) or without (Fig. 4E) sevoflurane. The ADL of slow PTNs was also unchanged by lightening anaesthesia.

### Latency, latency jitter, threshold and duration of antidromic responses

Figure 5 summarises the antidromic latency of 67 slow PTNs (Fig. 5A). ADLs were again measured from the first clear inflection in the antidromic spike in averages of responses to a single PT shock. Grey and black columns represent PTNs recorded respectively before (all 3 macaques) and after bicuculline injections (1 macaque). PTNs had latencies between 3.9 and 7.2 ms. This range would correspond to a range of conduction velocities of 12.4 m/s down to 6.6 m/s. Note the break in the abscissa between 0 and 4 ms: we intentionally avoided testing and optimising PTNs with shorter latencies, although many were recorded together with slow PTNs. In macaque O, there was no significant difference between ADLs of control PTNs (n=16) vs those recorded after bicuculline (n=23) (P>0.05, Wilcoxon rank sum test).

Slow PTNs showed little variation or jitter in ADL from one sweep to the next. We determined jitter by measuring the latency of each of around 100 consecutive sweeps with PT shocks at 1.2T. We had to exclude slow PTNs with very low signal-to noise and/or those with high spontaneous firing rates. For the 65 slow PTNs we could analyse, the jitter, defined as difference between earliest and latest antidromic spike within 2ms window around the mean antidromic spike, ranged from 86 µs to 917 µs (median 293 µs) for the first of two PT shocks given at ISI 10ms. Jitter was statistically significantly smaller for responses to the second shock (range 65 µs to 862 µs, median 187 µs, signtest p<0.001).

Figure 5B plots the antidromic thresholds of 61 slow PTNs for which the threshold could be determined. About half had thresholds of less than 300 µA. There was no significant correlation between ADL and threshold (n=61, r=-0.08, p>0.5).

Figure 5C plots the trough-to-peak duration of macaque slow PTNs. Unlike fast PTNs, many of which have ‘thin’ spikes (Vigneswaran et al 2011), it was noticeable that nearly all slow PTNs had rather broad spikes. For 55 well-isolated slow PTNs, we measured the duration of the spike from its negative trough to the succeeding positive peak (Vigneswaran et al. 2011). With one exception, slow PTN spikes had widths ranging from 0.39 to 1.1ms (mean 0.71ms, SD 0.14ms). The slow PTNs we sampled showed no obvious sign of initial segment-somadendritic (IS-SD) segmentation (Jankowska and Roberts 1972).

### Amplitude of PTN spikes

A major advantage of the probe used is that if it moves slightly, the recorded cell just moves to a different contact on the probe, rather than being lost completely. For all analyses we always used the channel with the largest peak-to-peak recorded spike; the amplitude was measured from the negative trough of the antidromic spike to either the preceding or succeeding maximum positivity, whichever was larger. Most of the slow PTNs showed great variability in spike amplitude across the different contacts. For example, the slow PTN shown with an asterisk in Fig. 6A had an ADL of 4.1 ms. The antidromic spikes recorded from each of the 32 contacts were averaged and these 32 averages have been superimposed in Fig. 6A. The spike had a peak-to-peak amplitude which varied from 104 µV on the middle contacts of the probe (see Fig. 6B) to a much smaller value on contacts located on the top and bottom of the probe. On these contacts spike amplitude fell below a pre-defined threshold, which was identified as 3 standard deviations above baseline activity, calculated from the level of activity between 21 and 24 ms after the onset of the second stimulus (i.e well after any antidromic or synaptic effects evoked by this stimulus). The size of the asterisks in Fig. 6B represent changes in relative spike amplitude across the contacts; the contacts with the smallest spikes are shown by grey asterisks.

**Figure 6.**
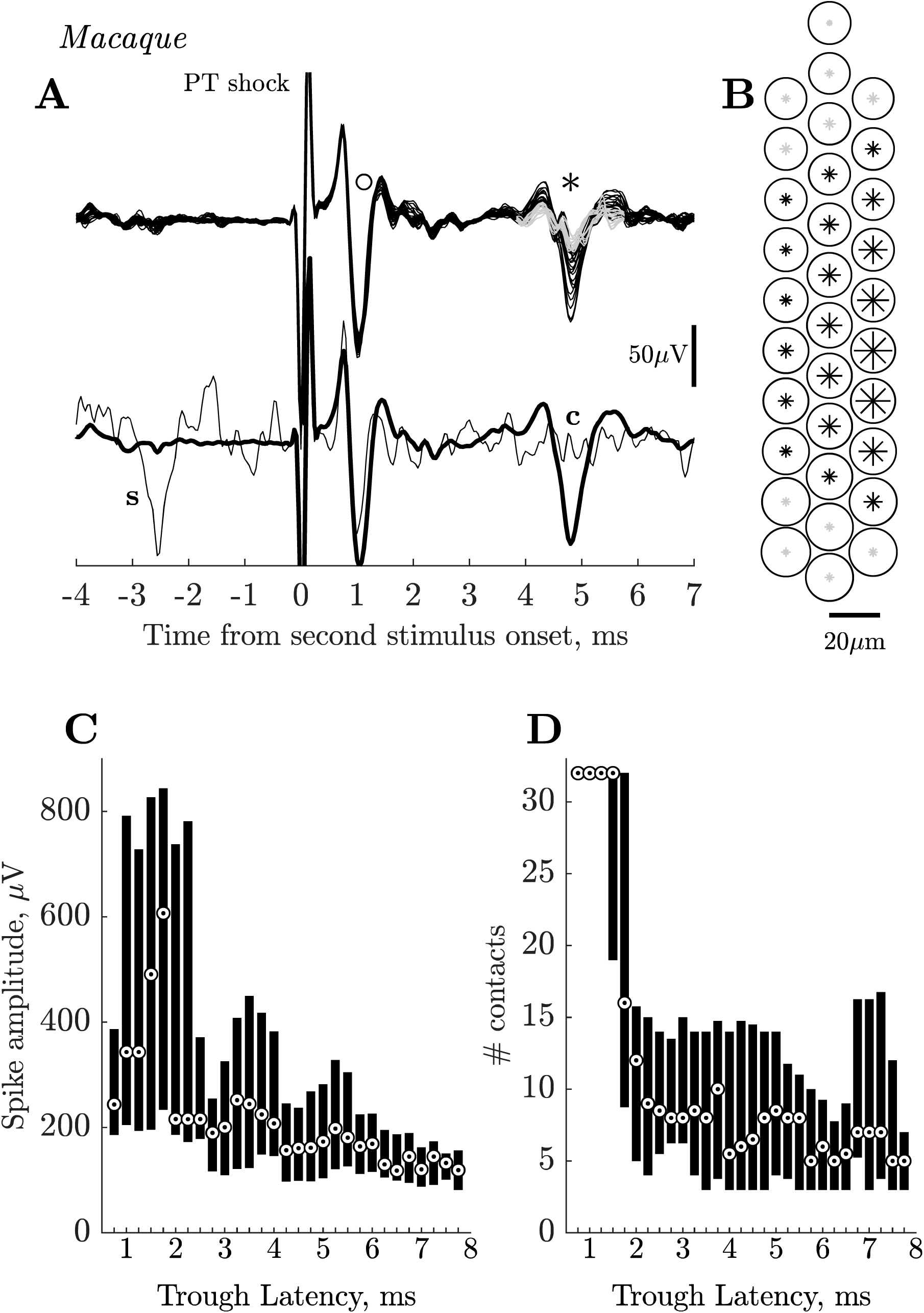
Macaque. Amplitude and spread of PTN spikes recorded on multiple contact silicon probe. **A.** Upper trace. Superimposed averages (each 100 sweeps) of antidromic activity simultaneously recorded on all 32 contacts of the polyprobe. The circle (o) marks a fast PTN with a short antidromic latency (< 1.0 ms), the asterisk (*) marks a slow PTN (ADL ∼ 4.0 ms). The early PTN had an almost identical amplitude on all 32 contacts, while the amplitude of the slow PTN varied significantly between contacts. Antidromic spikes of small amplitude (<∼20 µV) which did not cross a pre-defined threshold (see Results) are shown in grey. Lower traces show averages from the best contact for the slow PTN superimposed with a single sweep in which a spontaneous spike in the slow PTN (‘s’) collided the antidromic response in this PTN (‘c’). **B.** A map of polyprobe contacts. The size of circles (fast PTN) and asterisks (slow PTN) indicate the peak-to-peak amplitude of the corresponding antidromic spikes recorded across the array. Note the small variability in circle sizes (slightly larger towards the bottom of the probe) for fast PTN and much larger variability of asterisk sizes for slow PTN. Grey asterisks at the top and bottom of the probe correspond to the contacts where the size of the slow PTN was smaller than the estimated threshold. **C.** Median and 25-75 percentiles of the peak-to-peak spike amplitudes from PTNs with different negative trough antidromic latencies (measured from stimulus onset to the largest negative point on the recorded spike). Bin size is 1 ms sliding in 0.25ms steps, i.e. first bin includes data with peak latencies between 0.25 and 1.25 ms, second, between 0.5 and 1.5 ms etc. Circled dots indicate median values. Note that the depth of the recording probe was adjusted to give the maximum amplitude for all slow PTNs (> 4 ms) while this was not the case for all those with shorter ADLs, which happened to be present in the same recordings as the slow PTNs. Fast PTNs with early latencies (0.5-2 ms) had large amplitude spikes vs while slow PTNs with latencies > 4 ms had small spikes. These latencies correspond to conduction velocities ranging from ∼ 90 m/s for the fastest PTNs to ∼ 6m/s for the slowest. **D.** Median and 25-75 percentiles for the number of contacts on which antidromic spikes could be clearly distinguishable from noise. The possible maximum is 32 contacts; note that all fast PTNs with short ADLs were seen on all contacts.

By chance, at the same site at which this slow PTN was recorded, we also recorded a fast PTN with an early ADL of 0.7 ms (marked with a circle in Fig. 6A). This PTN had a spike amplitude (187 µV) which showed very little variation across contacts, as demonstrated by the near perfect overlap of the signals from all 32 contacts, and the similar size of the circles plotted in Fig. 6B to indicate action potential amplitude. The lower traces in Fig. 6A show the averages from the best contact for the slow PTN, and an example of collision of the slow PTN’s antidromic response.

We analysed all of the recordings to determine the distribution of amplitudes of PTN spikes across the different contacts. This analysis included the 67 slow PTNs with ADLs > 3.9 ms, but in addition other antidromic spikes that we could identify in the recording: these included 15 additional slow PTNs which we were not able to investigate fully because of their small amplitude, and 66 fast PTNs with ADLs from 0.7 to 3.9 ms which we had previously ignored because of our focus on slow PTNs. While we optimised the position of the probe for all of the 67 slow PTNs, we did not do this for any of these 81 additional slow and fast PTNs. Figure 6C shows the distribution of the peak-to-peak amplitudes of these spikes, for the contact that yielded the largest spike. The 25-75 percentile of the amplitudes, together with the median value, has been plotted against the ‘trough latency’ of the antidromic spike. This is the latency of the spike measured at its negative trough (for example, at the points indicated by ‘o’ and ‘*’ in Fig. 6A). This was easier to identify automatically than the onset latency (true ADL), especially for the smaller spikes or spikes contaminated by the field or other antidromic spikes.

Figure 6C shows that most of the faster PTNs with trough latencies shorter than 2.0 ms exhibited a wide variety of large spike amplitudes, even though we did not optimise probe position for these PTNs. These fast PTNs had median values rising from 200 µV to over 600 µV. In contrast, slow PTNs with ADLs > 4.0 ms were mostly small, with median values between 100 and 200 µV. There was a significant negative correlation between amplitude and latency of spike troughs when all the data in Fig. 6C were included (r=-0.42, p<0.001, Spearman). However, if only the slow PTNs with trough latencies > 4.25 ms were considered, the correlation disappeared (p> 0.2).

Figure 6D shows a corresponding analysis for the number of contacts upon which a given PTN could be clearly identified above the calculated threshold (see above). Due to contamination of the antidromic spike by other antidromic spikes or field responses, this analysis was possible for only 133 cells out of 148 used for amplitude analysis. The fastest PTNs were present on all 32 contacts, while slow PTNs were present on fewer contacts, with a median of between 5 and 9. These results suggest that the fastest PTNs make up a distinct subpopulation of M1 PTNs. There was a significant negative correlation between number of contacts and trough latency for all data in Fig. 6D (r=-0.37, p<0.001, Spearman). However, if only the slow PTNs with trough latencies > 4.25 ms were considered, the correlation disappeared (p> 0.7).

### Failure of antidromic responses

As stated above, every slow PTN tested with 3 x PT (ISI 3 ms) or 2 x PT (ISI 10 ms) shocks, at 1.2T, showed responses that followed high frequency stimulation. A few PTNs showed less than perfect following: for 2 x PT at 10 ms ISI, 5 of 57 tested PTNs (9%) showed a failure rate to the second shock of >10%, and the highest failure rate was 37% (74 failures out of 200 sweeps).

In the example shown in Fig. 7A-C, a PTN with an ADL of 5.9 ms and low antidromic threshold (85 μA), sometimes failed to respond to the second of two shocks at 1.2T (102 μA). On most sweeps (165/200; black traces in Fig. 7A) the antidromic response was present to the second shock, but in the remaining 35 (grey traces), the PTN failed to respond. There was a high level of spontaneous activity in other neurons included in this recording, but the failures were not due to collision of the antidromic response, since there were no spontaneous spikes immediately preceding the second shock (no spikes in the grey traces). On sweeps with failures after the second shock, failures also occurred after the first PT shock, although with a slightly lower rate (in the grey traces, some antidromic responses to the first shock were present). The averages of responses (black) and failures (grey) are shown in Fig. 7B. The failures showed no obvious clustering during the sequence of 200 sweeps: in Fig. 7C each vertical line indicates the occurrence of a sweep with a failed response to the second shock.

**Figure 7.**
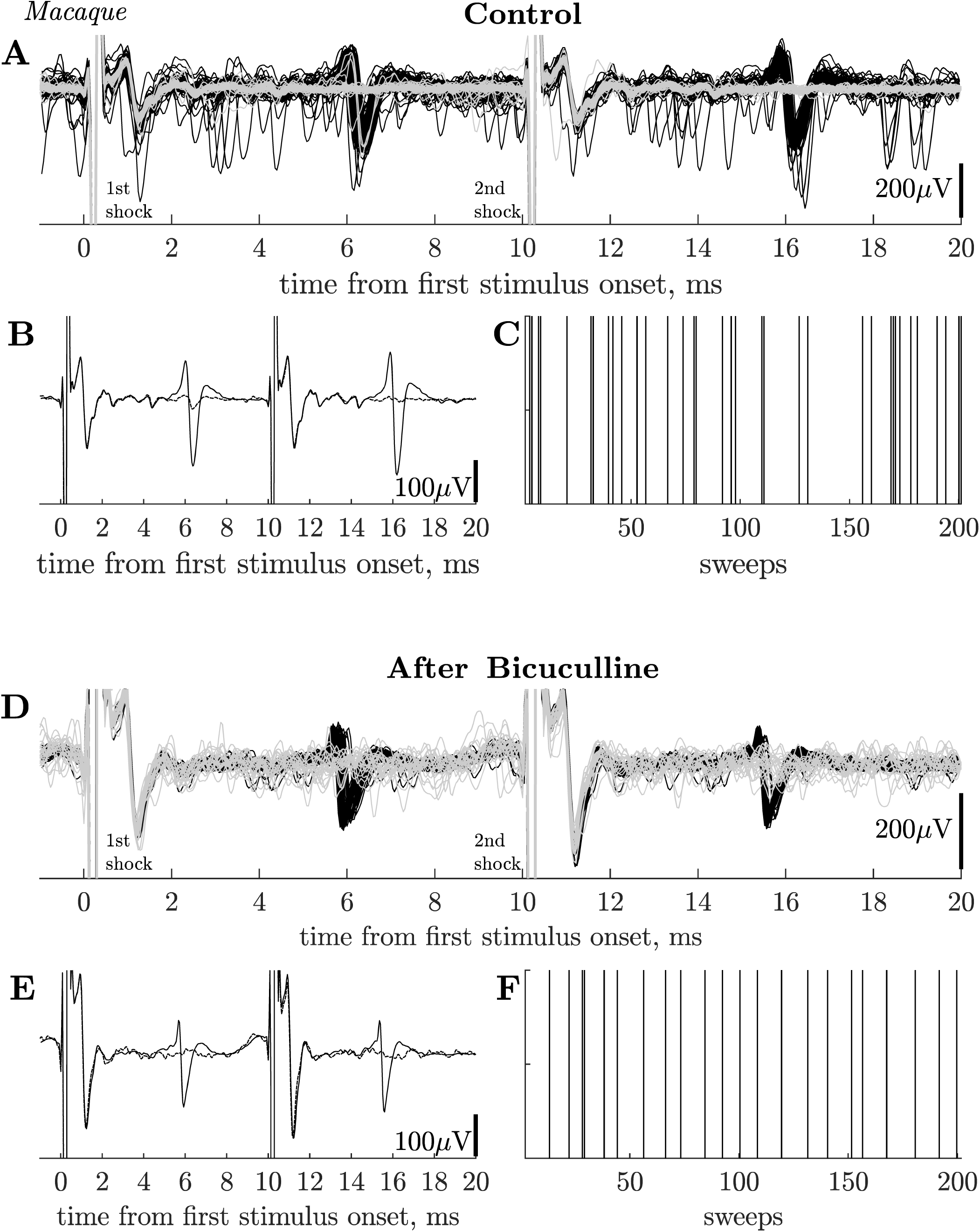
Macaque. Failure of antidromic invasion in slow PTNs. **A.** 200 superimposed sweeps showing responses of a slow PTN to paired PT shocks (ISI 10 ms) at 102 µA intensity. There were 35 failures of the antidromic response to the second shock (in grey). ADL of this PTN was 5.8 ms and threshold 85 µA. There was a high level of spontaneous activity in other neurons, but not in this PTN (no spontaneous spikes in grey sweeps and no collisions evident). Recordings taken in macaque O before bicuculline injection. **B.** Averaged sweeps for this PTN using all 165 sweeps in which the PTN responded to both shocks (black), and for 35 sweeps in which it failed to the second PT shock (dashed). **C.** Shows the timeline of the 200 paired stimuli; the vertical lines mark the sweeps on which the PTN failed to respond to the second shock. There was no obvious grouping of these failures. **D.** Recordings from monkey O from another PTN (ADL 5.5 ms and threshold 360 uA), recorded 59 min after an intracortical injection of bicuculline. In this case there were 22 failures in 200 sweeps (grey). Stimulus intensity 432 uA. **E.** Averaged sweeps for this PTN using all 178 sweeps in which the PTN responded to both shocks (black), and for 22 sweeps in which it failed to the second PT shock (dashed). **F.** as in C above.

Figure 7D-F shows a second case in which a slow PTN (ADL 5.5 ms) also showed regular failures (grey traces; 18 out of 200 sweeps). This PTN also failed to respond to both the first and second shocks on the same sweep. This PTN was recorded after an intracortical injection of bicuculline (see below). Once again there was no obvious clustering of the failure sweeps (Fig. 7F).

### Spinal termination of slow PTNs

In two macaques, we investigated whether slow PTNs could also be antidromically excited from the spinal cord, identifying them as corticospinal (example shown in Fig. 4B). Of 28 PTNs tested, 16 responded antidromically to shocks delivered to the spinal cord via electrodes at either C3 (12 PTNs) or C6 (4 PTNs).

### Effects of bicuculline

We looked for changes in the cortical responses to PT stimulation after bicuculline was microinjected into the motor cortex of two macaques. In monkey N, three injections (of 1, 1 and 5 µL of 50 µM bicuculline (25 µg/µL) were injected at around hourly intervals. The effects were very clear cut, so in the second experiment (macaque O) we tested a much lower concentration of 3 µM (1.5 µg/µL), making four 0.5 µL injections over a nine hour period, totalling 2 µL.

In both cases, within a few minutes of injection, bicuculline induced strong rhythmic bursting activity in M1, which was synchronous with seizure movements of the contralateral hand and digits. An example from macaque O is shown in Fig. 8A; these bursts, each with a duration of about 300-400 ms, occurred every few seconds. This bursting activity lasted for around one hour after injection.

**Figure 8.**
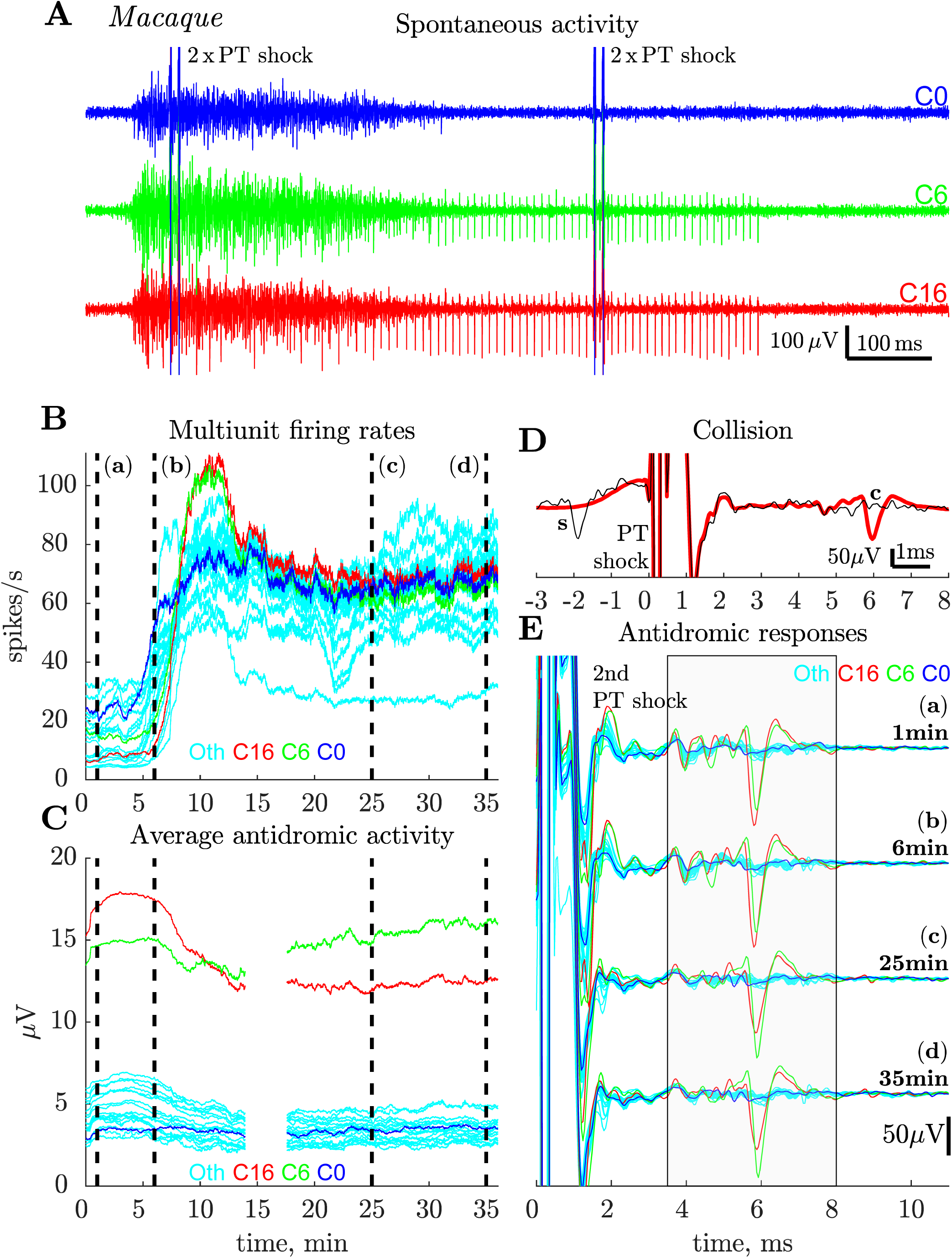
Macaque. Effects of intracortical bicuculline. **A.** Records showing spontaneous bursting of spike activity in M1 9.6 minutes after a microinjection of 0.5 µL of bicuculline (3µM, 1.5 µg/µL) close to the recording site. Each sweep is 1 s in duration and includes two consecutive pairs of PT stimuli (2 x PT) and contains one burst of activity lasting around 400 ms. Bursts of spikes in the cortex occurred about once per 2-3s and each burst was associated with strong seizures in the contralateral hand. Recordings from three different contacts (red, green and blue, C0, C6 and C16 respectively) on the polyprobe are shown; the same PTN (see D and E) was recorded on both the red and the green channels; it continued to fire regularly after the burst was over (300 to 700 ms). **B.** Plots showing frequency of multiunit spiking activity recorded in M1 just before the bicuculline injection was started (time zero), and then for 35 minutes after the injection. Multiunit activity was estimated on the same three channels as shown in A (same colour code) on which PTNs were recorded, and also from 21 other contacts (‘Oth’, light bue), with no PTNs present. Spikes were detected on each sweep, from 60 to 390 ms after PT stimulus onset (to exclude stimulation artefact and stimulus evoked activity), and averaged with a moving window of 100 s sliding with each sweep. Note the steep increase in firing rate on all contacts around 5 min after the onset of the injection, which after a peak around 10 min, maintained high rates. **C.** Plots of the averaged rectified activity that was recorded between 3.5 and 8.0 ms after the second of two PT shocks (ISI 10ms; intensity 750 µA) from 24 contacts (same colour code as in A and B) for the same part of the experiment as in B. This activity after the 2^nd^ shock was assumed to be antidromic, because, in general, synaptically-evoked activity did not follow two shocks. For each point activity was averaged with a moving window of 100 s (200 sweeps) sliding with each sweep (0.5s). Higher values of activity for two contacts (6 and 16, red and green) indicate presence of a slow PTN (ADL 5.7 ms) on these contacts. The activity in 22 other contacts (blue and light blue) remained fairly constant throughout the 35 min after injection onset, suggesting that no new PTNs were unmasked by bicuculline. Double shock recordings were not available for the period 14-17 min post-injection. **D.** Antidromic identification of the PTN recorded in A. Thick trace is average of 200 responses to a single shock at 750 µA. The single superimposed sweep shows a spontaneous spike in this PTN (‘s’) colliding (‘c’) the antidromic response. **E.** Responses to the second of two PT shocks (750 µA) at four different time points just after (a), 5 (b), 25 (c) and 35 (d) min after injection onset (see B). Before the recording, a slow PTN (ADL 5.7 ms, threshold 375 µA) was present on two contacts (6 and 16, red and green). It appeared slightly larger on the ‘red’ channel (a). The same PTN was present throughout the post-injection period (b-d) with slight changes in relative amplitude (larger on the green channel in (d), see also a switch in integrated antidromic activity red and green curves in (D), illustrating the same property). Although there were signs of several other small antidromic responses at 3-6 ms after the shock, there were no large increases or changes in these as a result of the bicuculline injection. Each curve is an average of 200 sweeps.

We also noticed a sharp increase in the overall spontaneous firing rate of multiunit activity at the layer V recording site (Fig. 8B). Multiunit firing rates were estimated from the number of threshold (median+ 4 std) crossings between 60 and 390 ms after the PT stimulus onset (to exclude any spikes in the period immediately following PT shocks). This included many of the periods of bursting such as that in the early part of Fig. 8A. We found that the spiking rate increased from around 5-20 spikes/s before injection to over 50 spikes/s and, for some channels, to over 100 spikes/s (red and green traces in Fig. 8A). These increases were seen on channels with antidromic PTN activity (red, blue, green) and on other channels with no signs of PTNs (cyan). Increases were sustained for at least 35 min after injection.

If antidromic spikes in slow PTNs were suppressed by recurrent inhibition, we would have expected a sustained increase in longer-latency antidromic activity after bicuculline. We assessed this activity by averaging rectified activity recorded on each channel for the period of 3.5-8 ms after the second of two PT shocks (intensity of each 450 µA) at ISI 10 ms. Since slow PTNs responded to both shocks, while other, synaptic effects failed to respond to the second shock, almost all of the activity after this second shock should be antidromic in nature. In contrast to spontaneous spiking activity, we saw no clear changes in the amount of slow antidromic activity evoked from the PT after bicuculline (Fig. 8C). Although antidromic activity was present on both the green and red probe channels (see Fig. 8A), these channels did not show any sustained increases after bicuculline. Other channels (blue, cyan) with less antidromic activity in the control period, also showed no increases after the injection. These results suggest that there was no additional late (3-8 ms) antidromic activity recruited as a result of the disinhibition generated by bicuculline.

Finally, we looked at antidromic responses in single neurons in the post-bicuculline period (Fig. 8E). One slow PTN (ADL 5.7 ms; identification and collision in Fig. 8D) with a large spike was present on two probe channels (red, green in Fig. 8A) both before (Fig. 8E (a)) and after the injection (b-d). Its ADL was unchanged by bicuculline injection. There was a slight drift in electrode position over the 35 min of recording, indicated by the change in relative spike size on these channels (red > green before and green> red after 35 min(d)), confirming that the electrode was still among PTNs. There were many small antidromic potentials on these and other channels (blue, multiple other channels represented by cyan traces), but these did not change as a result of the injection (cf Fig. 8C).

Similar results were obtained in the experiment in monkey N where a larger amount of bicuculline was injected. In this case there was again strong rhythmic bursting and seizures. Once again we saw increased spontaneous firing rates but no obvious changes in antidromic activity. These results demonstrate that injection of bicuculline did not reveal any additional slow PTNs.

In macaque O, we compared the characteristics of slow PTNs recorded before and after bicuculline injection. Figure 5 shows that the distributions of ADLs and antidromic thresholds were similar before (open bars) and after (shaded bars). We also assessed whether failure of antidromic invasion was changed by bicuculline. In this macaque, we tested 16 slow PTNs before bicuculline and 22 after; PTNs exhibiting signs of failure were actually not more common after bicuculline (6/22, 27%) than before (3/16, 19%, χ^2^ test p>0.5) for the 2 x PT paradigm, and the same was true for the 3 x PT paradigm (36% and 20%, respectively).

## Discussion

A full understanding of the corticospinal system depends upon the discovery of the function of all of its constituent fibres. While a great deal has been learned about its fast-conducting components, in humans and other animal species, our knowledge of the function of the slow fibres is rudimentary, despite the fact that they are well-defined in the neuroanatomical literature and make up the largest proportion of the corticospinal tract.

To study the function of these slow PTNs, it is essential to be able to record their activity and identify them electrophysiologically. We have demonstrated that it is possible to use a multiple contact silicon electrode to record from slow PTNs in both the rat and the macaque monkey. Our definition of ‘slow’ corresponds to an ADL of ∼4.0 ms or longer in the macaque, with an axonal conduction velocity of around 10 m/s or slower, and in the rat, ∼2.6 ms or longer in the rat, with again a conduction velocity of 10 m/s or slower. These slow PTNs were identified antidromically from the PT, and were characterised by the standard criteria of sharp threshold, relatively fixed latency with low jitter (especially after the second of two PT shocks), high frequency following, and, where possible, the collision test.

### Range of antidromic latencies of slow PTNs

The initial studies in the rat established that our recording methodology could yield PTNs with conduction velocities in the lowest range expected, with several PTNs having ADLs > 4 ms, longer than those reported in a previous study (Mediratta and Nicoll 1983). Using the same methodology we subsequently recorded slow PTNs from macaque M1 with ADLs ranging from 3.9 to 7.2 ms (Fig. 5; Fig. 9A, black bars). The sample in this study was clearly very different from that in many previous studies in the macaque, which have been dominated by PTNs with short ADLs (e.g. Evarts 1965; Humphrey and Corrie 1978; Ghosh and Porter 1988a; Firmin et al. 2014). Figure 9A compares the slow PTN ADLs recorded in this study (black bars) with those reported by Vigneswaran et al. (2011; grey bars). If we assume a conduction distance of 47 mm from PT to M1 (see Firmin et al. 2014), the slow PTNs sampled in these recordings came from PTNs with axon conduction velocities ranging from 12 down to 6.6 m/s, with a median value of 9.5 m/s. The relationship between conduction velocity and axon size is captured by the Hursh factor (Hursh 1939). If we assume a Hursh factor of 6 m/s for every micron of fibre diameter, the slowest conducting PTN which we were able to record would have had an axon with a diameter of just over 1 µm.

**Figure 9.**
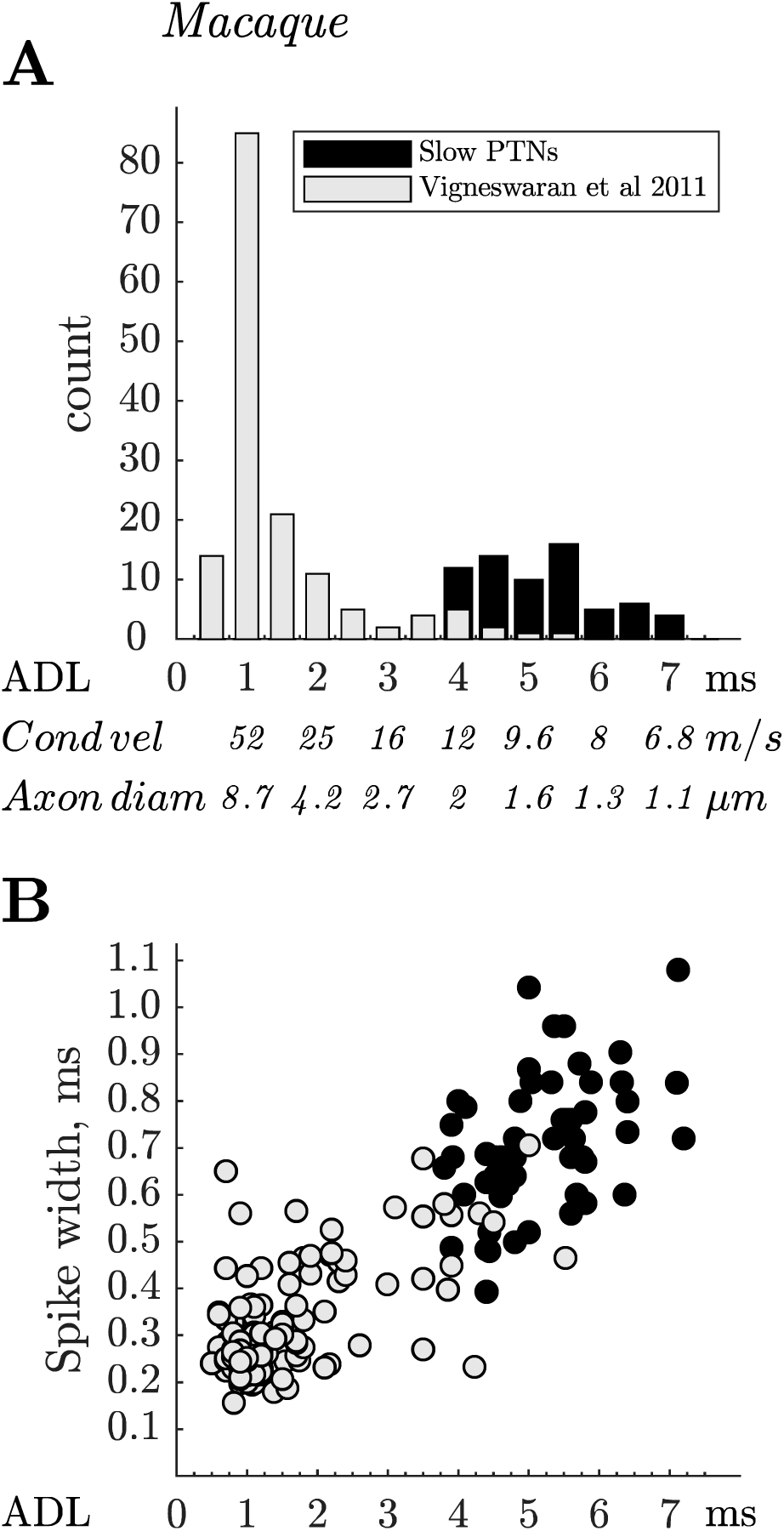
Macaque. Antidromic latencies and spike durations of fast and slow PTNs. **A.** Comparison of antidromic latencies of the 67 slow PTNs recorded from M1 in this study (black bars), with latencies of 151 M1 PTNs recorded two awake macaques using single platinum-in-glass microelectrodes (light grey bars; data from Vigneswaran et al. 2011). Note the very small numbers of slow PTNs in the latter study. **B.** Spike duration for the PTNs in this study (black circles, n= 55), plotted against the ADL of each PTN. Data from the Vigneswaran study is shown in grey circles. Note that the clear positive correlation between ADL and spike duration for fast PTNs extends into the slow PTN range, which typically had spike durations of 400 to 700 µS. The correlation between spike duration and ADL for the slow PTNs was significant (R=0.44, p<0.001).

In the rat, most ADLs ranged from 2.6 to 7.3 ms. An estimated conduction distance of 20 mm would correspond to conduction velocities of 7.7 m/s down to 2.7 m/s for the axons of these PTNs. Again, applying a Hursh factor of 6, the slower value would correspond to an axon diameter of around 0.5 µm. One exceptional PTN had an ADL of 14.6 ms (Fig. 1) which would have conducted at only 1.4 m/s with an estimated axon diameter of only 0.2 µm (if it were myelinated), similar to the lowest value reported by Leenen et al. (1985). The largest PT axons in the rat are around 3 µm and fastest conduction velocity is around 18 m/s (Mediratta and Nicoll 1983; Leenen et al. 1985).

It is known that in both macaque and rat, the PT as a whole contains many axons with diameters smaller than 1 µm (Häggqvist 1937; Leenen et al. 1985; Firmin et al. 2014) and a recent study by Innocenti et al. (2019) showed fine axons in both the PT and CST labelled by injections in macaque M1. The median value for M1 fibres in the PT were reported to be only 1.09 µm.

Thus although we report recordings from a population of slow PTNs, one might expect, on the basis of the anatomical literature, more slow PTNs with latencies longer than reported here. That discrepancy is particularly striking in the macaque, because, if assumptions about the Hursh factor are correct, none of the slow PTNs we recorded would have had axons with diameters smaller than 1 µm. We discuss below the different possible explanations for this finding.

### Are fine axons in the PT activated by test stimuli?

We employed a search stimulus of 750 µA with a biphasic configuration with each phase 100 µs in duration, which is the same as used in earlier studies of fine axons in the callosum (Waxman and Swadlow 1976) and corticospinal tract (Mediratta and Nicoll 1983). Most of the slow PTNs we recorded, in both macaque and rat, had thresholds of 300 µA or less (Figs. 2B and 5B, respectively), less than half the search stimulus intensity. We did routinely test higher strengths (up to 2 mA) to see if we had missed antidromic responses, and this yielded a few additional (4) PTNs, but certainly did not recruit a significant additional population of slower PTNs. Indeed, we found no significant relationship between ADL and antidromic threshold for slow PTNs (cf Firmin et al. (2014) for fast PTNs), and the slowest PTN we recorded (ADL 14.6 ms) had a threshold of only 75 µA (Fig. 1). We suggested previously that antidromic threshold shows a greater dependency on the location of a fibre relative to the stimulating electrode than on fibre size (Firmin et al. 2014). We conclude that the stimuli we used were effective in activating fine axons in the PT.

### Does electrode recording bias mean that slow PTNs are missed?

We decided to use the multiple contact silicon probe approach because slow PTNs are conspicuously lacking in the literature on PTNs in macaque primary motor cortex, and this may be partly due to the well-known bias, when using single metal extracellular microlectrodes, towards larger cells with faster axons (Towe and Harding 1970; Humphrey and Corrie 1978; Kraskov et al 2019). For example, in our earlier study of awake macaques (Firmin et al. 2014), recorded from M1 using metal microelectrodes, we found only 7% of PTNs with ADLs of > 4.0 ms. The longest ADL for M1 was 5.6 ms.

In contrast, the present study yielded many slow PTNs; the yield was 67 slow PTNs from 25 successful penetrations in the macaques. However, spikes from slow PTNs were generally small, difficult to isolate and unstable. Figure 6A and B show that spikes from slow PTNs exhibited considerable variation in amplitude across the 32 different contacts on the probe, and many were only discriminable on just a few contacts (Fig. 6D). In contrast, spikes from fast PTNs often had large amplitude spikes (Fig. 6A-C) and could be recorded on every contact on the probe (Fig. 6D). These properties suggest that the fastest PTNs represent a distinct subpopulation of cells with large somas and fast axons, consistent with the well-established fact that, when recording with single microlectrodes in M1, it is relatively easy to isolate fast PTNs, and to maintain stable recordings from them (see ‘Experimenter bias’ below).

Overall, we found significant negative correlations between ADL and spike amplitude on the optimal contact, and between ADL and the number of contacts on which the spike could be clearly recognised. But this correlation was primarily driven by the difference between fast and slow PTNs. The correlations for both measures disappeared if only slow PTNs with trough latency above 4.25 ms were taken into consideration (p>0.2 and p>0.7, respectively). Importantly, for the 67 slow PTNs (ADL 3.9-7.2 ms) whose amplitude was carefully optimised, there was no significant correlation between amplitude and ADL (p>0.15). This result suggests that soma size, which is probably the main factor influencing spike amplitude, and axon size (conduction velocity) may not be correlated for slow PTNs. Some studies which have compared PTN soma size with axon size have found a correlation (Sakai and Woody 1988) while others have not (Ghosh and Porter 1988a).

### Failure to evoke antidromic responses

Failure to evoke antidromic responses has been reported previously (Lipski 1981; Swadlow 1998). We saw some evidence of this phenomenon in the macaque recordings (Fig. 7). However, failure rates were generally low: for the 2 x PT shock test, only 9% of PTNs showed failure in > 10% of sweeps. Failures could occur both after the first and second shock of the 2 x PT pair (Fig. 7A). This limited degree of failure in the sampled PTNs would not have compromised their antidromic identification. These failures could have occurred because the axon in the PT was not activated (see above) or because the antidromic impulse failed to invade the cell body of the PTN. Failure to invade might be related to the degree of depolarisation at the moment that the antidromic spike arrives at the initial segment, but if so, it is remarkable that the number of PTNs showing some evidence of failure was not higher after bicuculline microinjections, which would have depolarised the PTNs, judging by their increased firing rate (Fig. 8B). Clearly failure would need to be much more severe to prevent identification as PTNs, but we cannot exclude that this might be a property of the very slowest PTNs and therefore explain their absence from our recordings.

The slow PTNs we sampled appeared to show only slight activity-dependent changes in ADL (Swadlow et al. 1978; Swadlow 1989). When a neuron discharges spontaneously, its axon undergoes periods of super-and subnormal conduction. These can lead to large changes (> 1 ms) in the latency of antidromic spikes in some systems, including callosal (Swadlow et al. 1978; Soteropoulos and Baker 2007) and bulbospinal neurons (Boers et al. 2005) but not others (Turner and Delong 2000). By using double shocks, it is possible to demonstrate that responses exhibiting latency jitter after the first shock show less jitter after the second shock, since those responses are less affected by preceding spontaneous activity (Swadlow et al. 1978).

Even though some of the slow PTNs in our sample were spontaneously active, they generally showed only small amounts of jitter (∼ 100 µs) in the ADL of responses to the first shock. As expected, jitter was usually even less after the second shock (see for example Fig. 1B (rat) and Fig. 7D (macaque)). These observations suggest that the super- and subnormal periods in slow PTN axons may be relatively short. They further suggest that it is unlikely that we missed any particularly slow PTNs because they had very variable ADLs and their responses were therefore not recognised as antidromic. Indeed, during recordings, we observed that whenever spikes showed responses with a highly variable latency to the first shock, they always failed to respond to the second shock, and we therefore classified such responses as synaptic in nature.

### Does recurrent inhibition block antidromic invasion of slow PTNs?

It has been suggested that one reason for the lack of slow PTNs in the electrophysiological literature is that PT stimulation excites the axons of fast PTNs with intracortical collaterals that could activate inhibitory inteneurons synapsing on slow PTNs (Innocenti et al. 2019). As a result, the fast recurrent inhibition of these PTNs would begin before the arrival of antidromic potentials from their axons, with the consequence that antidromic invasion might be prevented.

Recurrent inhibition (RI) has been reported in a number of studies using intracellular recording from pyramidal neurons (Phillips 1959; Suzuki and Tukahara 1963; Stefanis and Jasper 1964; Takahashi et al. 1967), all which show that RI is long-lasting and frequency dependent. The onset latency of RI in pyramidal neurons after PT stimulation has been reported to be between 8 and 40 ms. Ghosh and Porter (1988b) made intracellular recordings from a small number of slow PTNs in macaque motor cortex and found that recurrent IPSPs could begin as early as 1.5-5.0 ms after a single PT shock, and last for 40-50 ms. Interestingly, these IPSPs did not prevent antidromic activation and identification of these slowly-conducting neurons. If RI explains blockage of antidromic responses, we anticipated that it would be more pronounced for the second of two paired PT shocks (10 ms ISI) and that failure would generally be higher for repetitive than for single stimuli.

We did not find any evidence of this sort in our sample of slow PTNs. All of them showed following of both 3 x PT, with an ISI of 3 ms, and 2 x PT at 10 ms ISI, despite the fact that later shocks should have sent antidromic volleys into slow PTNs at times when RI should have been most pronounced. In PTNs showing some failures, these occurred both on single shocks, and to a similar extent for both the first and second shocks of the 2 x PT pair (Fig. 7).

For PTNs with low antidromic thresholds, one might argue that RI from fast PTNs might be weak because only small numbers of their fibres would be activated by weak stimuli. We explored this by applying large shocks (up to 2 mA) to see if this would block any of the antidromic responses (e.g. Fig 4E). This was not the case in the 41 macaque PTNs that we tested with such strong stimulation.

Since RI is likely to be reduced by anaesthesia, we compared antidromic responses with and without supplementary gas anaesthesia, but again never saw any significant depression of antidromic responses when anaesthesia was lightened.

### Attempts to block recurrent inhibition by intracortical microinjections of bicuculline

Since recurrent inhibition is known to be mediated by GABA_A_ inhibitory synapses, it has been suggested that blocking GABA_A_ action might reveal slow PTNs whose antidromic responses are blocked by RI. This was tested in two macaques. In the first monkey, we tried small volumes of a 50 µM solution injected close to the M1 recording site, and this produced strong bursting activity in the cortex and seizure activity in contralateral hand and digits. In the second monkey we found that a much weaker concentration (3 µM or 1.5 µg/µL) produced very similar effects. This latter, lower dose is at the low end of a range of concentrations used in previous studies in the monkey (Matsumura et al. 1991, 1995; Wang et al. 2000; Galineau et al. 2017). It was still very effective in inducing bursting activity and seizures (Fig. 8A) and in significantly increasing firing rate at the nearby recording site (Fig. 8B). However despite these striking effects, bicuculline, at either the high or low concentrations used, did not unmask any new antidromic effects, either at the antidromic mass potential (Fig. 8C) or single unit level (Fig. 8E), despite the fact that the recording probe was maintained in layer V throughout the post-injection period, as evidenced by the continued presence of slow PTNs recorded prior to injection.

Two further points are worth making: in one macaque we compared the ADLs and antidromic thresholds of PTNs recorded before and after bicuculline, and these were very similar (Fig. 5A, B). In particular, we saw no evidence for PTNs with longer ADLs (> 8 ms) after bicuculline. The second point is that if antidromic failure is the result of RI, one might predict a lower rate of failure after bicuculline. But in the few PTNs that did show some failures (Fig. 7D-F), this was not the case.

It could also be argued that recurrent facilitation from other corticospinal axons could interfere with antidromic activation of slow PTNs (Kraskov et al. 2019). However, given that bicuculline would be expected to increase these effects greatly (by blocking any antagonistic inhibitory effects), and that we did see greatly increased spontaneous activity, it is significant that antidromic responses were unaffected.

In conclusion, although one might expect RI to be reduced by the GABA_A_-blocker bicuculline, this did not reveal new populations of slow PTNs.

### An additional factor: experimenter bias

There is also an element of experimenter bias to consider as a further explanation of the dearth of slow PTNs in published recordings. Because recordings from fast PTNs are generally larger in amplitude and more stable than those from smaller, slow unstable PTNs, there is a tendency not to pursue slower PTNs. This may be particularly true in awake animals performing a trained task, where it is essential to retain recordings from the same neuron throughout acquisition of sufficient behavioural trials to allow full analysis. In the current study, we would probably not have collected as many slow PTNs had we not, at the outset, deliberately excluded focussing on the fast PTNs that were present in the recordings. The use of more sensitive recording probes makes it possible to partly offset the bias towards fast PTNs, and these methods will be needed if we are to gain insights into the activity of slow PTNs during well-characterised motor and other behaviours. We are also aware that some slow PTNs, although antidromically activated, may be ‘silent’, not showing any spontaneous, task-related discharge, as has been documented in the macaque callosal system (Soteropoulos and Baker 2007).

### Sampling the very slowest PTNs

In the macaque recordings, we did not find any slow PTNs with ADLs longer than 7.2 ms (estimated CV 6.6 m/s). One possibility is that spikes generated by slower-conducting PTNs, with ADLs > 8 ms, may have been just too small to record, even with the silicon probe. However, as pointed out above, there was no obvious correlation between ADL and spike amplitude for slow PTNs, so it is not obvious why these slowest PTNs went undetected, unless they represent another distinct neurophysiological sub-population.

Given the fact that the macaque PT contains many fibres with diameters of < 1 µm, we might have expected that these very slow PTNs would be present in greater numbers. Applying a Hursh factor of 6, the very smallest axons (∼ 0.2 to 0.5 µm; Innocenti et al. 2019) emanating from M1 should conduct with velocities as low as 1-2 m/s axons, which would give an estimated ADL > 20 ms.

However, even when we searched in the recordings for additional very small but identifiable antidromic responses, we never found any beyond 8 ms (see Fig. 6C and D) and antidromic activity of any kind was not apparent at these long latencies (see Fig. 8E, for example). Thus an alternative explanation might be that the Hursh factor of 6 does not apply to fine myelinated axons in the macaque corticospinal tract. This has never been tested directly. Fine fibres may display other differences from fast fibres which could influence conduction velocity, including the ratio of myelin to fibre diameter and the inter-nodal distance (Waxman and Swadlow 1977; Ritchie 1982; Innocenti et al. 2019). So the possibility remains that if the Hursh factor were higher for fine axons (e.g. 12-15 rather than 6), the predicted minimum conduction velocities would be much closer to those we sampled (i.e. ∼ 6 m/s). This would explain while we never saw signs of antidromic activity representing even lower CVs.

### Prospect: functional properties of slow PTNs

To date, our knowledge of corticospinal function appears to have been informed mostly by the physiology of the fast-conducting components (Firmin et al. 2014). Understanding the slow-conducting axons of the corticospinal tract is important for at least two reasons: their large numbers in relation to the fast PTNs, and the fact that they may play a significant role in recovery from damage to the CST, since they are less vulnerable to trauma than the fast fibres (Blight 1991; Quencer et al. 1992), and it is known that trauma can change myelination and conduction properties (Sampaio-Baptista and Johansen-Berg 2017).

The present study already revealed some potentially important features of slow PTNs which might give some clues to their function. Firstly, we showed that some of these slow PTNs project into the spinal cord. It is known that corticospinal projections from M1 target mostly the intermediate zone of the spinal grey matter and the ventral horn, largely avoiding the dorsal horn laminae (Ralston and Ralston 1985; Morecraft et al. 2013). If there were a general relationship between PTN soma size and axon diameter (Sakai and Woody 1988), then it is interesting to note that some of the layer V corticospinal neurons that were transneuronally labelled by (Rathelot and Strick 2006) had very small soma diameters, hinting that some slow PTNs could make direct, cortico-motoneuronal connections. Monosynaptic effects from these PTNs would be expected to have late onsets, and such effects have been reported (Maier et al. 1998; Witham et al. 2016).

The long conduction times in slow PTNs from cortex to cervical spinal cord, which we estimate to be around 15 to 20 ms for those sampled here (vs ∼ 4-5ms for faster PTNs), would seem to preclude them from contributing to the initiation of fast upper limb movements or to rapid transcortical reflexes involving CM cells (Cheney and Fetz 1984; Scott 2004). However slow PTNs might contribute to slower changes in postural set or reflex excitability (Tanji et al. 1978).

Figure 9B shows another interesting property of macaque slow PTNs. They have broad spikes, with a trough-to-peak duration of 0.4 to 0.7 ms (Fig. 9B, black circles). This contrasts strongly with the ‘thin’ spikes exhibited by fast conducting PTNs in M1 (Fig. 9B grey circles, taken from the dataset of (Vigneswaran et al. 2011; see also Takahashi 1955, for the first observation of this kind, made on cat PTNs). For the slow PTN group reported here there was strong positive correlation between spike duration and ADL (r=0.44, p=0.001). The brief spike durations in fast PTNs may be related to their capacity to discharge high frequency bursts of action potentials; it will be important to discover the discharge patterns of slow PTNs in awake animals, and whether they follow rules that may govern the structure and function of neurons with fine axons (see Perge et al. 2012). As mentioned above, the low jitter in ADL possibly hints at fast recovery cycles in slow PTNs.

In terms of synaptic inputs to slow PTNs, these will need to be determined in the awake animal and compared with the rich proprioceptive inputs known to influence fast PTNs in M1 (Lemon and Porter 1976; Wong et al. 1978; Lemon 1981). The original proposal that slow PTNs receive strong RI from fast PTNs may need to be revisited.

## Conclusions

We have demonstrated that it is possible to record from slow PTNs in M1 of both macaque and rat using a multiple contact silicon probe. Thus we have partly addressed the well-known discrepancy in the literature between the distribution of PT axon diameters and antidromic latencies/conduction velocities. The results suggest that the main factor is the bias towards recording from fast PTNs, which represent a distinct sub-population of M1 PTNs. Although we could isolate spikes from slow PTNs with the probe, they were generally small and difficult to record for long periods. Importantly, we found no evidence that recurrent inhibition of slow PTNs by fast conducting PT fibres is a factor in masking or blocking antidromic invasion. Failure of antidromic impulses to invade slow PTNs was only rarely found. Whether or not we sampled the entire range of slow PTNs in M1 remains open, but we have at least shown that it is possible to record from some of these neurons. This study therefore opens up the possibility of investigating them in the awake animal and thereby advancing our understanding of the contributions made by the small and numerous fine fibres within the PT (and possibly in other central pathways) in both health and disease.

## Acknowledgements

This work was supported by grants from BBSRC (DSS) BB/P019757/1 and MRC (SNB) MR/P012922/1 to SN Baker and Wellcome Trust (AK). The authors would like to thank Dr Kathy Murphy, Dr Rocio Palacios-O’Connor and Dr Chris Blau for assistance with anaesthesia, and Norman Charlton, Terri Jackson and Andrew Atkinson for technical support. We are grateful to Peter Kirkwood for his comments on the manuscript.

## References

1. Blight AR. 1991. Morphometric analysis of a model of spinal cord injury in guinea pigs, with behavioral evidence of delayed secondary pathology. J Neurol Sci. 103:156–171.

2. Boers J, Ford TW, Holstege G, Kirkwood PA (2005). Functional heterogeneity among neurons in the nucleus retroambiguus with lumbosacral projections in female cats. J. Neurophysiol. 94: 2617–2629.

3. Cheney PD, Fetz EE. 1984. Corticomotoneuronal cells contribute to long-latency stretch reflexes in the rhesus monkey. J Physiol (Lond). 349:249–272.

4. Collins GF, Baker SN. 2014. Automated intracellular recording with multiple sharp micropipettes. Presented at the Society for Neuroscience Annual Meeting. Washington, DC: SfN.

5. Evarts EV. 1965. Relation of discharge frequency to conduction velocity in pyramidal tract neurons. J Neurophysiol. 28:216–228.

6. Firmin L, Field P, Maier MA, Kraskov A, Kirkwood PA, Nakajima K, Lemon RN, Glickstein M. 2014. Axon diameters and conduction velocities in the macaque pyramidal tract. J Neurophysiol. 112:1229–1240.

7. Galineau L, Kas A, Worbe Y, Chaigneau M, Herard A-S, Guillermier M, Delzescaux T, Féger J, Hantraye P, Tremblay L. 2017. Cortical areas involved in behavioral expression of external pallidum dysfunctions: A PET imaging study in non-human primates. Neuroimage. 146:1025–1037.

8. Ghosh S, Porter R. 1988a. Morphology of pyramidal neurones in monkey motor cortex and the synaptic actions of their intracortical axon collaterals. J Physiol (Lond). 400:593–615.

9. Ghosh S, Porter R. 1988b. Corticocortical synaptic influences on morphologically identified pyramidal neurones in the motor cortex of the monkey. J Physiol (Lond). 400:617–629.

10. Häggqvist G. 1937. Faseranalytische studien über die pyramidebahn. Acta Psychiatr Scand. 12:457–466.

11. Humphrey DR, Corrie WS. 1978. Properties of pyramidal tract neuron system within a functionally defined subregion of primate motor cortex. J Neurophysiol. 41:216–243.

12. Hursh JB. 1939. Conduction velocity and diameter of nerve fibers. American Journal of Physiology-Legacy Content. 127:131–139.

13. Innocenti GM, Caminiti R, Rouiller EM, Knott G, Dyrby TB, Descoteaux M, Thiran J-P. 2019. Diversity of cortico-descending projections: histological and diffusion MRI characterization in the monkey. Cereb Cortex. 29:788–801.

14. Jankowska E, Roberts WJ. 1972. An electrophysiological demonstration of the axonal projections ofsingle spinal interneurones in the cat. J Physiol (Lond). 222:597–622.

15. Kraskov A, Dancause N, Quallo MM, Shepherd S, Lemon RN (2009). Corticospinal neurons in macaque ventral premotor cortex with mirror properties: a potential mechanism for action suppression? Neuron 64: 922–930.

16. Kraskov A, Baker S, Soteropoulos D, Kirkwood P, Lemon R. 2019. The corticospinal discrepancy: where are all the slow pyramidal tract neurons? Cereb Cortex. 29:3977–3981.

17. Kuypers H. 1981. Anatomy of the descending pathways. In: Brookhart JM, Mountcastle VB, editors. Handbook of Physiology - The Nervous System II. Bethesda, MD: American Physiological Society. p. 597–666.

18. Lassek AM. 1941. The pyramidal tract of the monkeys. A Betz cell and pyramidal tract enumeration. J Comp Neurol. 74:193–202.

19. Leenen LP, Meek J, Posthuma PR, Nieuwenhuys R. 1985. A detailed morphometrical analysis of the pyramidal tract of the rat. Brain Res. 359:65–80.

20. Lemon RN. 1981. Functional properties of monkey motor cortex neurones receiving afferent input from the hand and fingers. J Physiol (Lond). 311:497–519.

21. Lemon RN. 1984. Methods for neuronal recording in conscious animals. John Wiley & Sons.

22. Lemon RN. 2008. Descending pathways in motor control. Annu Rev Neurosci. 31:195–218.

23. Lemon RN, Porter R. 1976. Afferent input to movement-related precentral neurons in conscious monkeys. Proc Biol Sci. 194:313–339.

24. Lipski J. 1981. Antidromic activation of neurones as an analytic tool in the study of the central nervous system. J Neurosci Methods. 4:1–32.

25. Macpherson J, Wiesendanger M, Marangoz C, Miles TS. 1982. Corticospinal neurones of the supplementary motor area of monkeys. A single unit study. Exp Brain Res. 48:81–88.

26. Maier MA, Illert M, Kirkwood PA, Nielsen J, Lemon RN. 1998. Does a C3-C4 propriospinal system transmit corticospinal excitation in the primate? An investigation in the macaque monkey. J Physiol (Lond). 511 (Pt 1):191–212.

27. Matsumura M, Sawaguchi T, Oishi T, Ueki K, Kubota K. 1991. Behavioral deficits induced by local injection of bicuculline and muscimol into the primate motor and premotor cortex. J Neurophysiol. 65:1542–1553.

28. Matsumura M, Tremblay L, Richard H, Filion M. 1995. Activity of pallidal neurons in the monkey during dyskinesia induced by injection of bicuculline in the external pallidum. Neuroscience. 65:59–70.

29. Mediratta NK, Nicoll JA. 1983. Conduction velocities of corticospinal axons in the rat studied by recording cortical antidromic responses. J Physiol (Lond). 336:545–561.

30. Morecraft RJ, Ge J, Stilwell-Morecraft KS, McNeal DW, Pizzimenti MA, Darling WG. 2013. Terminal distribution of the corticospinal projection from the hand/arm region of the primary motor cortex to the cervical enlargement in rhesus monkey. J Comp Neurol. 521:4205–4235.

31. Perge JA, Niven JE, Mugnaini E, Balasubramanian V, Sterling P. 2012. Why do axons differ in caliber? J Neurosci. 32:626–638.

32. Phillips CG. 1959. Actions of antidromic pyramidal volleys on single Betz cells in the cat. Q J Exp Physiol Cogn Med Sci. 44:1–25.

33. Quallo MM, Kraskov A, Lemon RN (2012). The activity of primary motor cortex corticospinal neurons during tool use by macaque monkeys. J Neurosci 32:17351–17364.

34. Quencer RM, Bunge RP, Egnor M, Green BA, Puckett W, Naidich TP, Post MJ, Norenberg M. 1992. Acute traumatic central cord syndrome: MRI-pathological correlations. Neuroradiology. 34:85–94.

35. Ralston DD, Ralston HJ. 1985. The terminations of corticospinal tract axons in the macaque monkey. J Comp Neurol. 242:325–337.

36. Rathelot J-A, Strick PL. 2006. Muscle representation in the macaque motor cortex: an anatomical perspective. Proc Natl Acad Sci USA. 103:8257–8262.

37. Ritchie JM. 1982. On the relation between fibre diameter and conduction velocity in myelinated nerve fibres. Proc R Soc Lond, B, Biol Sci. 217:29–35.

38. Russell JR, Demyer W. 1961. The quantitative corticoid origin of pyramidal axons of Macaca rhesus. With some remarks on the slow rate of axolysis. Neurology. 11:96–108.

39. Sakai H, Woody CD. 1988. Relationships between axonal diameter, soma size, and axonal conduction velocity of HRP-filled, pyramidal tract cells of awake cats. Brain Res. 460:1–7.

40. Sampaio-Baptista C, Johansen-Berg H. 2017. White matter plasticity in the adult brain. Neuron. 96:1239–1251.

41. Scott SH. 2004. Optimal feedback control and the neural basis of volitional motor control. Nat Rev Neurosci. 5:532–546.

42. Soteropoulos DS, Baker SN. 2007. Different contributions of the corpus callosum and cerebellum to motor coordination in monkey. J Neurophysiol. 98:2962–2973.

43. Stefanis C, Jasper H. 1964. Recurrent collateral inhibition in pyramidal tract neurons. J Neurophysiol. 27:855–877.

44. Suzuki H, Tukahara Y. 1963. Recurrent inhibition of the Betz cell. Jpn J Physiol. 13:386–398.

45. Swadlow HA, Waxman SG, Rosene DL (1978) Latency variability and the identification of antidromically activated neurons in the mammalian brain. Exp Brain Res 32:439–443.

46. Swadlow HA. 1989. Efferent neurons and suspected interneurons in S-1 vibrissa cortex of the awake rabbit: receptive fields and axonal properties. J Neurophysiol. 62:288–308.

47. Swadlow HA. 1998. Neocortical efferent neurons with very slowly conducting axons: strategies for reliable antidromic identification. J Neurosci Methods. 79:131–141.

48. Takahashi K (1965) Slow and fast groups of pyramidal tract cells and their respective membrane properties. J Neurophysiol 28: 908–924.

49. Takahashi K, Kubota K, Uno M. 1967. Recurrent facilitation in cat pyramidal tract cells. J Neurophysiol. 30:22–34.

50. Tanji J, Taniguchi J, Fukushima K (1978) Relation of slowly conducting pyramidal tract neurons to specific aspects of forearm movement. J de Physiol (Paris) 74: 293–296.

51. Towe AL, Harding GW. 1970. Extracellular microelectrode sampling bias. Exp Neurol. 29:366–381.

52. Turner RS, Delong MR (2000) Corticostriatal activity in primary motor cortex of the macaque. J Neurosci 20: 7096–7108.

53. Vigneswaran G, Kraskov A, Lemon RN. 2011. Large identified pyramidal cells in macaque motor and premotor cortex exhibit “thin spikes”: implications for cell type classification. J Neurosci. 31:14235–14242.

54. Wang M, Zhang H, Li B-M. 2000. Deficit in conditional visuomotor learning by local infusion of bicuculline into the ventral prefrontal cortex in monkeys. European Journal of Neuroscience. 12:3787–3796.

55. Waxman SG, Swadlow HA. 1976. Morphology and physiology of visual callosal axons: evidence for a supernormal period in central myelinated axons. Brain Res. 113:179–187.

56. Waxman SG, Swadlow HA. 1977. The conduction properties of axons in central white matter. Prog Neurobiol. 8:297–324.

57. Witham CL, Fisher KM, Edgley SA, Baker SN. 2016. Corticospinal Inputs to primate motoneurons innervating the forelimb from two divisions of primary motor cortex and area 3a. J Neurosci. 36:2605–2616.

58. Wong YC, Kwan HC, MacKay WA, Murphy JT. 1978. Spatial organization of precentral cortex in awake primates. I. Somatosensory inputs. J Neurophysiol. 41:1107–1119.

